# The novel shapeshifting bacterial phylum *Saltatorellota*

**DOI:** 10.1101/817700

**Authors:** Sandra Wiegand, Mareike Jogler, Timo Kohn, Ram Prasad Awal, Sonja Oberbeckmann, Katharina Kesy, Olga Jeske, Peter Schumann, Stijn H. Peeters, Nicolai Kallscheuer, Mike Strauss, Anja Heuer, Mike S. M. Jetten, Matthias Labrenz, Manfred Rohde, Christian Boedeker, Harald Engelhardt, Dirk Schüler, Christian Jogler

## Abstract

Our current understanding of a free-living bacterium - capable of withstanding a variety of environmental stresses-is represented by the image of a peptidoglycan-armored rigid casket. The making and breaking of peptidoglycan greatly determines cell shape. The cytoplasmic membrane follows this shape, pressed towards the cell wall by turgor pressure. Consequently, bacteria are morphologically static organisms, in contrast to eukaryotic cells that can facilitate shape changes. Here we report the discovery of the novel bacterial phylum *Saltatorellota*, that challenges this concept of a bacterial cell. Members of this phylum can change their shape, are capable of amoeba-like locomotion and trunk-formation through the creation of extensive pseudopodia-like structures. Two independent *Saltatorellota* cells can fuse, and they employ various forms of cell division from budding to canonical binary fission. Despite their polymorphisms, members of the *Saltatorellota* do possess a peptidoglycan cell wall. Their genomes encode flagella and type IV pili as well as a bacterial actin homolog, the ‘saltatorellin’. This protein is most similar to MamK, a dynamic filament-forming protein, that aligns and segregates magnetosome organelles via treadmilling. We found saltatorellin to form filaments in both, *E. coli* and *Magnetospirillum gryphiswaldense*, leading to the hypothesis that shapeshifting and pseudopodia formation might be driven by treadmilling of saltatorellin.

## Introduction

The bacterial *Planctomycetes*-*Verrucomicrobia*-*Chlamydiae* (PVC) superphylum comprises extraordinary bacterial phyla of medical, environmental and biotechnological importance [1, 2]. Among those, the phylum *Chlamydiae* is most intensely studied. It consists primarily of pathogens with small, reduced genomes that divide without the canonical bacterial cell division protein FtsZ [3]. The presence or absence of peptidoglycan (PG) was debated for decades - a controversy known as the chlamydial anomaly [4–9]. Today, the presence of PG, which is required for cell division, is accepted [7, 9–12]. Not all *Chlamydiae* are pathogens, some might even be symbionts known as environmental *Chlamydiae* [13]. However, this term is misleading as all described members of the phylum *Chlamydiae* are obligate intracellular bacteria [13]. The phylum *Verrucomicrobia* comprises free-living species with unparalleled cell shapes exemplified by *Verrucomicrobium spinosum* [14]. *Akkermansia muciniphila* [15] and other species that reside in the human intestinal tract can act as probiotics [16], influence obesity [17–19] or cancer treatment [20]. Other members of the phylum display unique metabolic traits such as thermoacidophilic methanotrophy [21, 22] or performance of the Knallgas reaction [23]. Similar to *Chlamydiae*, the presence of peptidoglycan in all phylogenetic branches of the phylum was only recently determined [24]. However, and in contrast to *Chlamydiae*, species of the phylum *Verrucomicrobia* encode FtsZ [25].

Members of the phylum *Planctomycetes* are ubiquitous bacteria that play major roles in global carbon and nitrogen cycles [2]. They perform the unique anammox reaction [26] by converting ammonium to dinitrogen gas [27]. Many *Planctomycetes* dwell on all sorts of marine surfaces [28–33] where they can dominate biofilms [34] and digest complex carbon substrates [35, 36]. Like *Chlamydiae* [10] and *Verrucomicrobia* [24], *Planctomycetes* were only recently found to possess peptidoglycan [37, 38]. With exception of *Candidatus* Brocadiaceae [39, 40], the anammox Planctomycetes, planctomycetal internal structures were rather found to be invaginations of the cytoplasmic membrane [41–43] than intracytoplasmic compartments as stated before [41, 44, 45]. *Planctomycetes* divide without FtsZ [3, 46, 47], either by polar, lateral or arbitrary budding [47, 48], or by binary fission.

In a recent study, we brought many novel planctomycetal strains into axenic culture and compared their characteristics [47]. The isolates of a phylogenetically deep branching group have rather small genomes compared to their peers and they are particularly rich in extracytoplasmic function σ factors (ECFs) [47]. They are most unique, as the cells divide by polar budding and binary fission [47]. Such exceptions from canonical *Planctomycetes* motivated us to investigate these strains further. Here we show how members of this group can shift their shape, divide in multiple different ways and are capable of amoeba-like locomotion. We reveal indications for cell fusions and the formation of trunk-like protuberances. We exclude osmotic artefacts by determining the strains natural habitat and analyse their genomes towards their unusual traits. Finally, we validly describe the three novel species *Saltatorellus ferox*, *Engelhardtia mirabilis* and *Rohdeia mirabilis* and propose that they do not belong to the phylum *Planctomycetes* but are affiliated to the novel phylum *Saltatorellota* within the PVC superphylum.

## Results and Discussion

During the last years, we isolated and characterized 78 novel planctomycetal strains [47]. Among them were three deep branching strains that appeared quite unusual in terms of cell division: cells of *Saltatorellus ferox* Poly30^T^ were shown to proliferate by binary fission or by budding [47]. To study this extraordinary behavior in greater detail, we employed time-lapse light microscopy and observed that *Engelhardtia mirabilis* Pla133^T^ cells also seem to be dividing by budding and binary fission (Fig. 1a and Movie S1). In addition, some cells divide only after massive swelling and it appears that in these cells, the cytoplasmic membrane divides first, only later followed by the outer membrane (Fig. 1a and Movie S1).

**Fig. 1.**
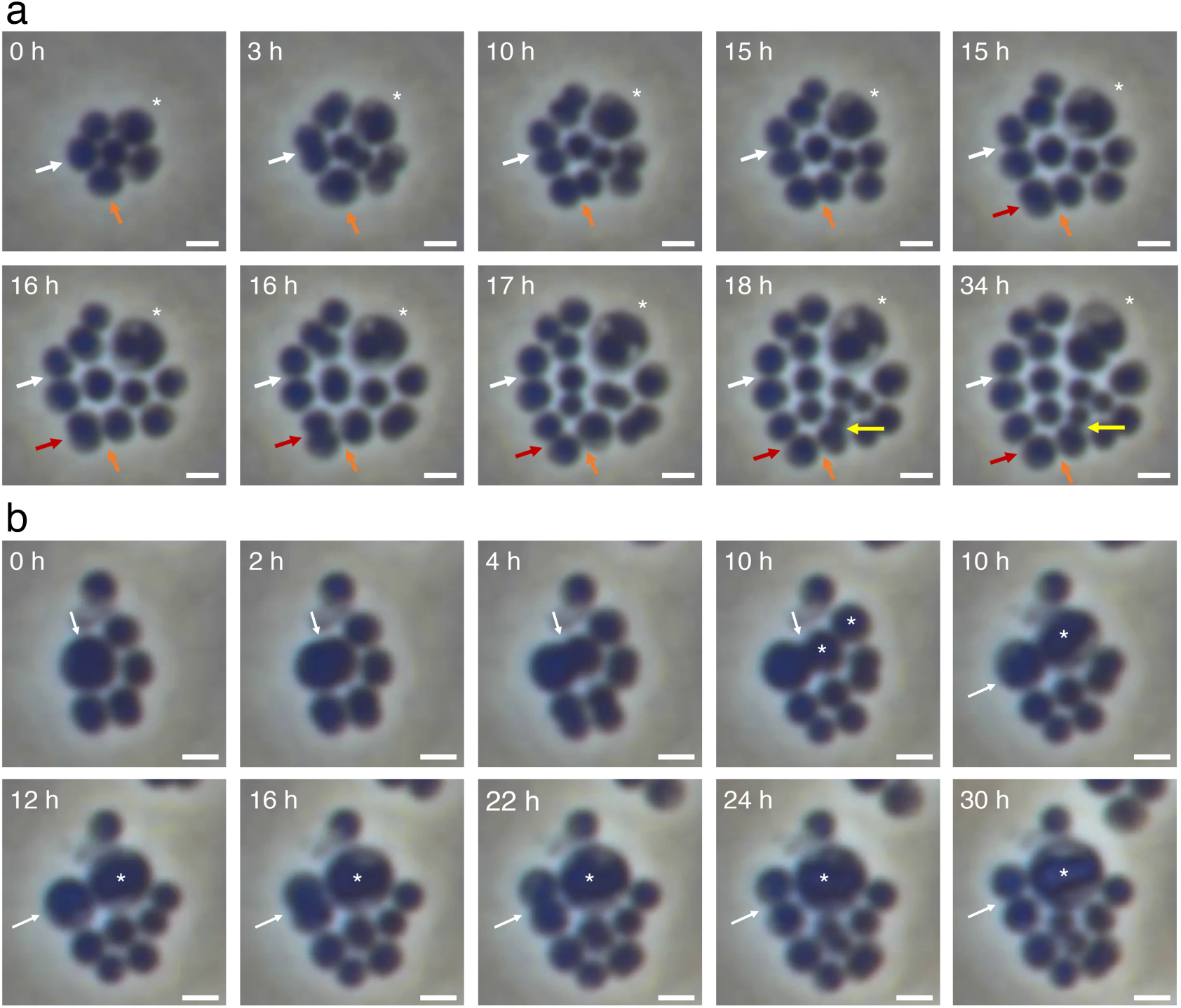
*Engelhardtia mirabilis* Pla133^T^ – division and fusion observed with time-lapse microscopy. **a)** Cells of *E. mirabilis* Pla133^T^ divide by multiple different mechanisms. Cells divide by binary fission (white, red and orange arrows) or polar budding (yellow arrow). Some cells (asterisk) swell and seem to divide first on the inside by a sort of budding where the outermost membrane follows the division process only later (34 h). **b)** Cells of *E. mirabilis* Pla133^T^ can fuse. A cell starts to divide by a sort of asymmetric binary fission (0-4 h, white arrow), when the daughter cell suddenly fuses with an adjacent cell (10 h, asterisk). The resulting cell (12-30 h, asterisk) appears to reshape the cytoplasm until it seems to divide again like the giant cell in a) (18-34h, asterisk). In contrast, the mother cell divides again by then symmetric binary fission (10-30 h, white arrow). Scale bar 1 µm.

Even more unusual, *E. mirabilis* Pla133^T^ cells appear to be fusing (Fig. 1b and Movie S2). To our knowledge such a cell fusion was never before observed among bacteria. However, it is almost impossible to determine if the fusing cells were entirely separated before, or if some kind of membrane-enclosed channel, maybe a remainder of an incomplete earlier division, facilitated the cell fusion. This is partly because acknowledged techniques to visualize such structures - as for example FM4-64 membrane staining - were not operational for *Saltatorellota* cells in our hands.

However, such unusual and unique interactions were not limited to *E. mirabilis* Pla133^T^, and once we started to re-analyze *S. ferox* Poly30^T^, we got even more intriguing results. In Fig. 2a and Movie S3 we were able to capture an incomplete and reversed budding process, followed by cell division by binary fission. Later, the daughter cell was found moving while changing shape. This shapeshifting seems to be key to the locomotion process and is reminiscent of the amoeboid locomotion of protists. Cells in Fig. 1 and 2a were immobilized by an agar cushion, a process well established for analyzing planctomycetal cells in our laboratory preventing flagella-mediated movements. If single moving cells were analyzed at higher magnification and temporal resolution, it became evident that the shape of *S. ferox* Poly30^T^ is changing in order to cause the amoeba-like locomotion. Parts of the cell seem to reach out for a new direction (Fig. 2b and Movie S4, 0-35 min), similar to pseudopodia of amoeba. Then, the entire cell appears to be pulled to the new location quite fast (Fig. 2b and Movie S4, 35-40 min). This behavior seems to be occurring in a small fraction of newly formed daughter cells, maybe implying some kind of cell differentiation (Movie S5).

**Fig. 2.**
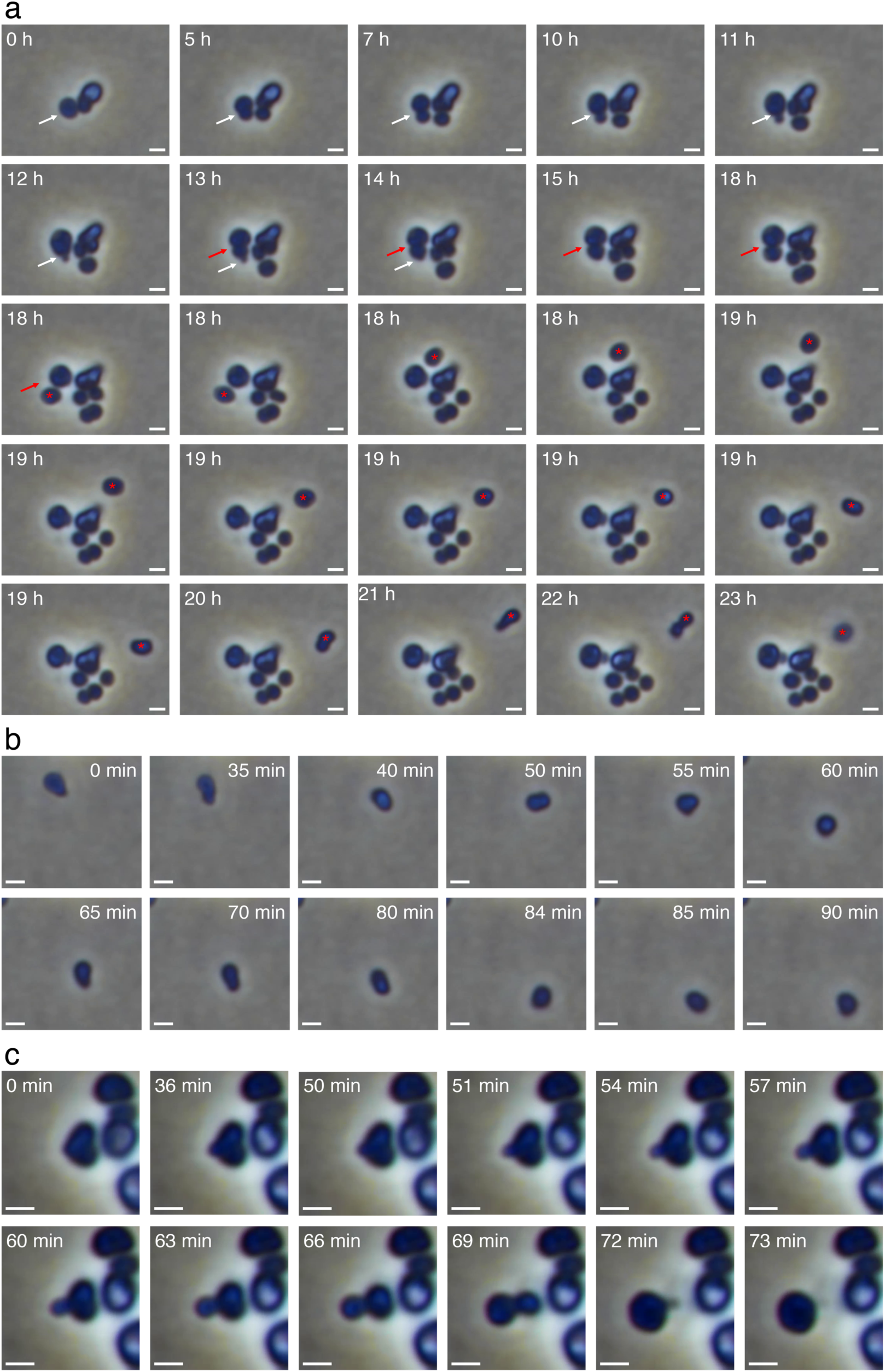
Locomotion and shapeshifting of *Saltatorellus ferox* Poly30^T^ observed with time-lapse microscopy. **a)** A cell seems to divide by budding (0-10 h, white arrow), but the budding process appears to be reversed (11-12 h, white arrow) while a second division by asymmetric binary fission occurs afterwards (13-18 h, red arrow). Once the daughter cell is separated (18 h, red asterisk), it starts to move (19-23 h) while locomotion is related with shapeshifting. **b)** At higher temporal and spatial resolution, a single cell moves despite the agar cushion by changing its shape. It reminds of amoeboid movement. **c)** A heart-shaped cell of strain Poly30^T^ (0 min) seems to start dividing by budding (0-60 min). The putative mother cell shrinks (63-69 min) and fuses again with the putative daughter cell (72-73 min). This way, via partial budding and subsequent refusion the cell moved on the agar cushion. Scale bars 1 µm.

Potential artefacts such as depressions in the agar cushions that allow cells to float by Brown motion could be excluded not only by careful specimen preparation, but by observation: if cells cross the focal plane, their shape appears blurry and ring like artefacts appear around them (moving cells in Movie S5). However, cells shapeshift and move in the focal plane as well (Movie S5).

A similar process intertwined with intense shapeshifting, although without ongoing progressive motion, becomes most obvious in Fig. 2c and Movie S6. The heart-shaped cell on the left-hand side seems to start a budding division cycle, but then just tunnels to its new position, where it adopts a spherical shape. Afterwards, the cell starts a successful reproductive cycle and releases three consecutive daughter cells by budding. Most of the described motility traits were observed if individual cells, or cells that were part of small aggregates were analyzed. However, if larger aggregates were explored, an additional type of protuberance formation became visible: the formation of a trunk-like structure, a structure formed and retracted like an amoeboid pseudopodium grasping for food particles (Fig. 3 and Movie S7). Formation and retraction of the structure took about 50 minutes. In our experimental setting, other than shapeshifting, trunk-formation did not result in any locomotion. However, both phenomena do parallel the formation of pseudopodia by amoeba that either sense the environment to initiate an endocytic or pinocytotic event or lead to locomotion.

**Fig. 3.**
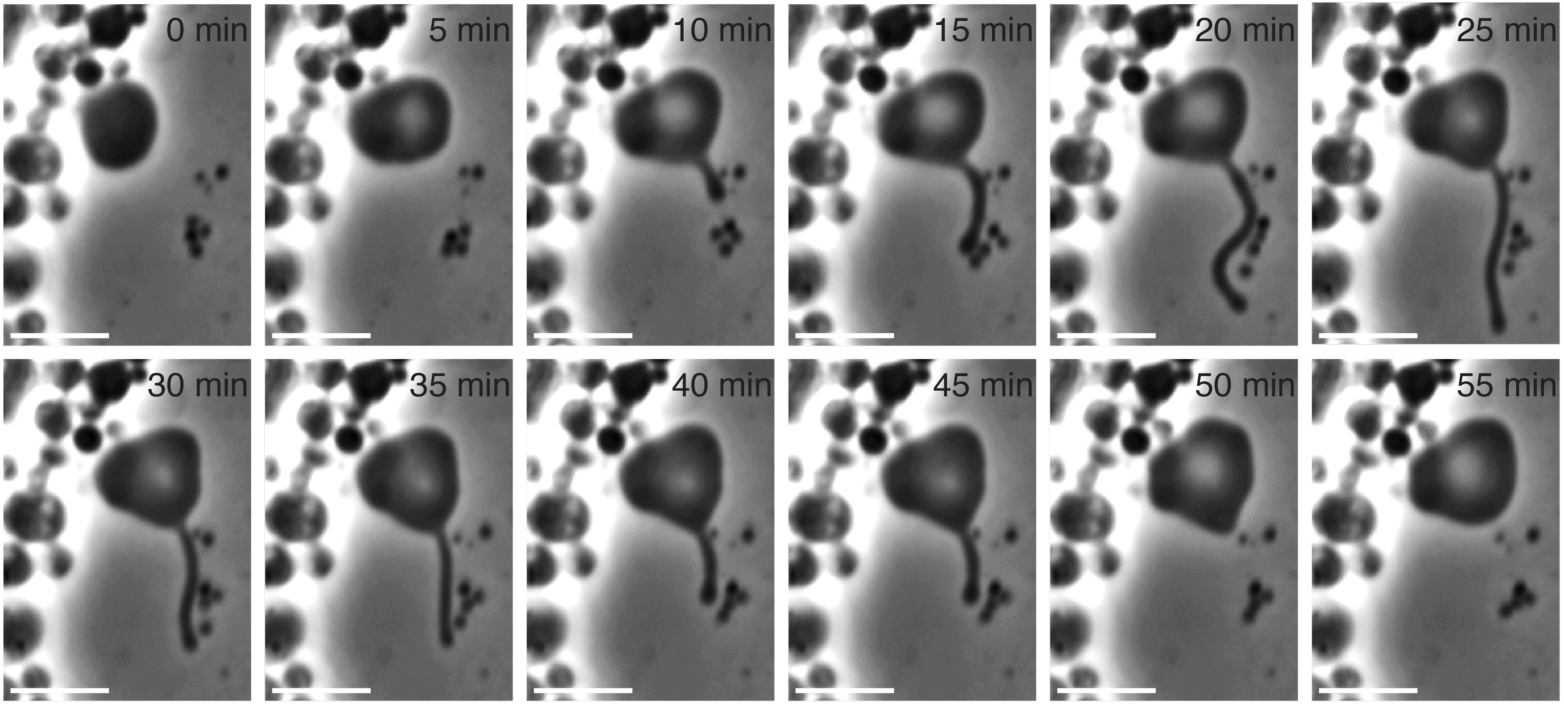
Trunk formation of a giant *Saltatorellus ferox* Poly30^T^ cell observed with time-lapse microscopy. In particular at the edge of huge aggregates, formed by hundreds of cells, single giant Poly30^T^ cells appear (Fig. S1) that can form trunk-like protuberances. Such structures remind of eukaryotic amoeboid pseudopodia for food absorption. Scale bar 5 µm.

To complement these light microscopic observations, we also employed different electron microscopic methods. Using scanning electron tomography (SEM), we were able to reproduce the different ways of cell division, showing that cells from the same axenic culture can divide either by budding, or by binary fission (Fig. 4a). Additionally, we employed cryo-electron tomography (CET) and analyzed frozen-hydrated cells of *E. mirabilis* Pla133^T^. The cell depicted in Fig. 4b divides by binary fission, just like a canonical coccoid Gram-negative bacterium. Going through the four different planes, multiple interconnected cytoplasmic invaginations become visible. Although such enlargements of the periplasmic space and tubular-vascular networks where previously described for *Planctomycetes* [41, 42], *E. mirabilis* Pla133^T^ appears quite unusually, more detailed and with a higher degree of curved membranes, compared with cells of the phylum *Planctomycetes*.

**Fig. 4.**
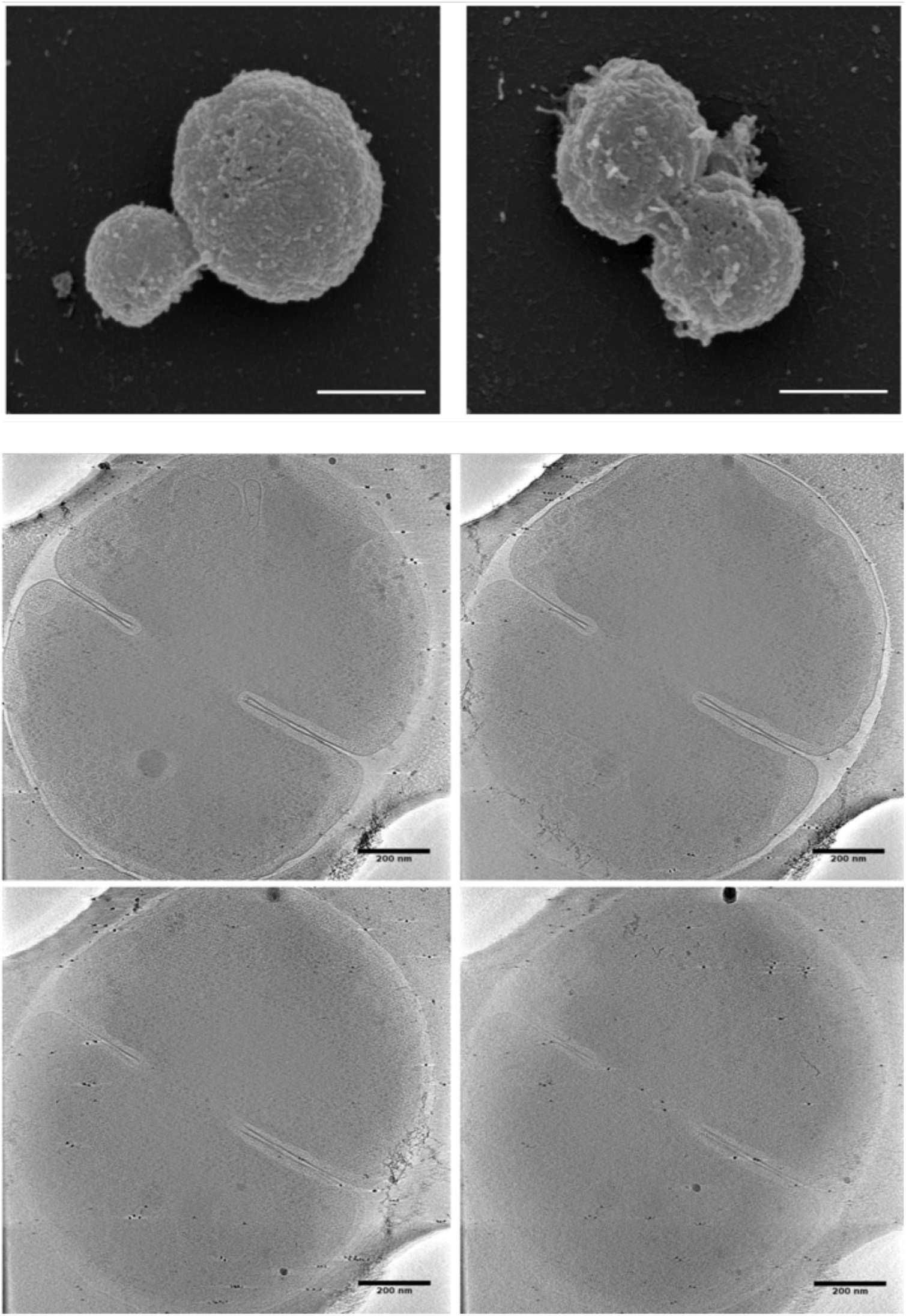
*Saltatorellota* divide by binary fission and budding. **a)** *Saltatorellus ferox* Poly30^T^ divides either by budding or by binary fission. The left and the right SEM micrograph show cells of the same species that divides by budding and binary fission, respectively. Scale bar 500 nm. **b)** Cryo-electron tomography of an *Engelhardtia mirabilis* Pla133^T^ cell. Four slices of the tomogram are shown that illustrate typical septum formation during binary fission of a coccoid cell. The cell is very flat and has multiple invaginations of the cytoplasmic membrane. Scale bar 200 nm.

A bigger picture of the various, above described traits emerges when approaching not only individual cells, but when focusing on the large aggregates they usually form when axenically cultured (Fig. S1). In growing aggregates (Movies S8 and S9) especially such traits as shapeshifting, motility and trunk formation can be observed. In Movie S10, a distinct trunk and a single motile cell appear in close proximity. Together with Movie S5 where at least one daughter cell from each microcolony moves away from the mother cell, one might speculate on a scouting function of this kind of differentiated cells. Precise quantification of the different phenotypes was hampered by this aggregate formation as even smaller aggregates span so many focal planes that only one layer can be observed at a time.

Dissolved solutes within bacteria cause significant osmotic pressure that is counteracted by peptidoglycan (PG), a rigid polymeric mesh of linear glycans [49] cross-linked by short peptides [50]. Among all free-living bacteria, PG is essential for the maintenance of cell integrity [24, 38]. Therefore, the unusual cell biology of *Saltatorellota* should be prevented by the rather static peptidoglycan (PG) sacculus [51].The synthesis of PG, together with FtsZ and 11 other canonical proteins, is essential for septal formation during cell division [52]. Thus, most bacteria comprise both, a PG cell wall and FtsZ. However, all three *Saltatorellota* strains were found to lack FtsZ along with most other canonical cell division proteins. Thus, while PG has been found in all PVC phyla, we wondered if members of the *Saltatorellota* might be different in this regard. This question was further urged by the observation that *Saltatorellota* cells are rather fragile. Centrifugation (4000 g, 10 min) seems to inactivate a significant proportion of cells and even careful blotting of electron microscopic grids on a paper tissue appeared to destroy the outer membrane of some cells. Furthermore, their general behavior reminds of cell wall-deficient L-form bacteria [53–55], or members of the PG-lacking phylum *Tenericutes, e.g. Mycoplasma* [56], that show comparable changes in shape, but are osmotically fragile and live host-associated [56].

To address the question of PG presence, we employed a methodology that was used before to detect PG in *Planctomycetes* [38] and *Verrucomicrobia* [24]. Firstly, we analyzed *Saltatorellota* cells for the presence of diaminopimelic acid (DAP). This non-proteinogenic amino acid is the diagnostic trifunctional linker in most Gram-negative PG sacculi and can be detected by characteristic mass over charge peaks in GC-MS spectra. *E. mirabilis* Pla133^T^ showed clear DAP-specific signals (380, 324, 306 and 278 m/z), while *S. ferox* Poly30^T^ showed only weak signals at 380, 324 and 278 m/z (Fig. S2). Nevertheless, the presence of PG is at least very likely in *E. mirabilis* Pla133^T^ and *S. ferox* Poly30^T^. However, a continuous sacculus cannot be concluded from DAP analysis. In Fig. 4b - where the PG layer of *E. mirabilis* Pla133^T^ is generally visible - it appears slightly discontinuous. This phenomenon is well known, caused by the method-inherent missing wedge as previously observed in e.g. *Planctomycetes* [38]. For *Rohdeia mirabilis* Pla163^T^, none of the DAP-specific signals could be detected at 380m/z and 306 m/z. Therefore, we prepared PG sacculi from this strain and TEM analysis clearly showed continuous PG sacculi as typical for Gram-negative bacteria (Fig. S3). *R. mirabilis* Pla163^T^ forms an extensive extracellular matrix, creating almost tissue-like aggregates. Given that DAP detection is based on wet cell mass the extensive matrix might have pushed cell numbers below DAP detection limit.

Taken together, all isolated *Saltatorellota* strains apparently possess a continuous PG sacculus that might undergo intensive modulations during shapeshifting, amoeba-like locomotion or trunk-formation.

All novel *Saltatorellota* isolates were obtained from microplastic particles incubated for two weeks either in the Baltic Sea or an inflowing river estuary with salinities of 1-1.5% as described before [47, 57]. In order to exclude osmotic stress and to gain deeper insights into the natural habitat of these strains, water samples were taken along the southern and eastern coastline of the Baltic Sea. We measured a salinity gradient of 4-9 PSU (approximately 0.4- 0.9%) along the transect (Fig. 5). *Saltatorellota* strains accounted for nearly 2% of the bacterial community in particle-attached fractions of the samples but were sparse in free-living fractions (Fig. 5, Table S1). Thus, *Saltatorellota* should be able to cope at least with salinity shifts between 0.4 and 1.5% under environmental conditions. To determine the optimal salinity for our *Saltatorellota* isolates, we tested media compositions with differing artificial seawater concentrations and observed optimal growth at concentrations corresponding to salinities of approximately 1-3%. Thus, we performed all our time-lapse experiments at two different salinities, both providing optimal growth conditions (2.03% and 1.35%), without noting a difference in cell behavior.

**Fig. 5.**
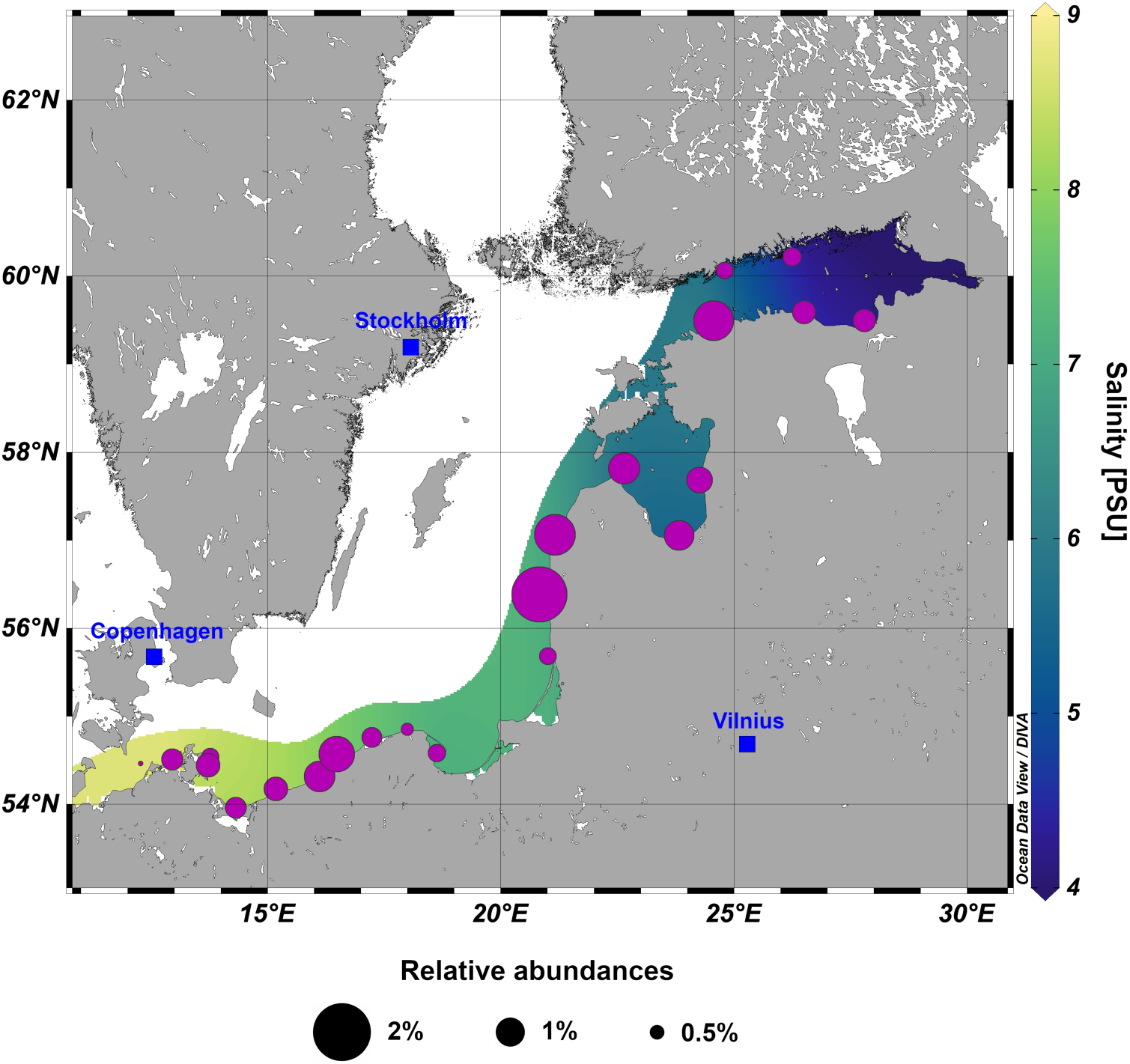
Natural occurrence of *Saltatorellota* strains in northern European Baltic Sea. The mean relative abundances (indicated by dot sizes) of 16S rRNA gene sequences from the particle-attached fraction determined by >97% identity to *Saltatorellota* strains Pla133^T^, Pla163^T^ and Poly30^T^. The background color scheme gives the salinity range along the southern and eastern coastline.

Consequently, our observations such as shapeshifting cannot be explained by osmotic artefacts. Furthermore, we conclude that the *Saltatorellota* strains are not osmotically fragile, despite their need to constantly re-modulate their peptidoglycan layer.

Since neither absence of PG, nor osmotic fragility could explain the unusual behavior of the *Saltatorellota* strains, we were wondering if they might encode proteins responsible for bacterial motility such as swimming, gliding, twitching or sliding [58] (Table S2). All three *Saltatorellota* isolates harbor the gene set required for building a canonical bacterial flagellum, localized in two mostly conserved clusters at distinct loci (Fig. S4, Table S2 and S3). From a genomic perspective, all three strains should therefore be capable to form a bacterial flagellum. Unexpectedly, *S. ferox* Poly30^T^ and *E. mirabilis* Pla133^T^ did not do so under the tested conditions. For >300 cells on 20 micrographs we did not observe a single flagellum. Neither did any of our light-microscopic observations or time-lapse series indicate any flagella-propelled swimming activity for both strains. However, in three TEM micrographs of *R. mirabilis* Pla163^T^, a flagellum was visible, but never attached to the cell body. Therefore, the observed locomotion is most likely unrelated to flagellar formation.

Besides flagella, we identified a second type of bacterial motility genes in the three *Saltatorellota* strains: genes encoding proteins for twitching motility (Fig. S5, Table S2 and S3). Key to this type of locomotion are long, thin appendages called type IV pili (T4P) [59]. These T4Ps are polymerized and depolymerized to fulfil their functions, that besides twitching motility are the uptake of DNA, biofilm or microcolony self-organization and many more [59]. T4P-mediated movement of the cell results from cycles of T4P elongation, surface adhesion and retraction [59]. T4Ps are thereby used as ‘grappling hooks’ leading to a characteristic jerky movement of cells [60]. However, what we observed for our *Saltatorellota* strains is very unlike this typical twitching movement. The closest relatives of *Saltatorellota*, the *Planctomycetes*, were also found to possess T4Ps, but twitching motility was never observed among this phylum [61]. Accordingly, they seem to lack the ATPase required for filament retraction. In contrast, *Saltatorellota* strains do encode PilB and PilT (Fig. S5, Table S2 and S3), the ATPases required for T4P formation and retraction. Thus, the T4P system might be involved in *Saltatorellota* locomotion or trunk-formation, but in an unseen manifestation. Furthermore, it could as well be responsible for the almost tissue-like microcolony and biofilm.

Nonetheless, canonical twitching motility cannot explain the observed amoeba-like locomotion or shapeshifting capability of *Saltatorellota* cells. In amoeboid cells such traits require a complex interplay of cytoskeletal elements (actin) with motor proteins (myosin). We were able to exclude the presence of motor proteins in the *Saltatorellota* isolates by mining the *Saltatorellota* genomes for homologs of eukaryotic myosin, kinesin and dynamin. On the other hand, the *Saltatorellota* strains each harbor two actin homologs [47]. In a detailed cluster analysis (see Material and Methods), we determined the position of these homologs in relation to all putative actin homologs in the BLAST nr database. We found that all three *Saltatorellota* strains encode MreB, the canonical actin homolog in bacteria, and a novel actin homolog that we suggest naming ‘saltatorellin’ (Fig. 6a). Saltatorellin branches close to MamK, an actin-like filament found in magnetotactic bacteria [62] such as *Magnetospirillum gryphiswaldense* (Fig. S6, Table S4). MamK was found to provide the sole driving force for magnetosome movement by pole-to-midcell treadmilling growth of cytomotive actin-like filaments [63]. Thus, we fused the saltatorellin gene of *S. ferox* Poly30^T^ to mCherry and found the product of this fusion construct to form filaments in *E. coli* and *M. gryphiswaldense* cells (Fig. 6b,). Our findings open the possibility that saltatorellin has properties comparable to MamK and might drive shapeshifting and amoeba-like locomotion in *Saltatorellota* by treadmilling. However, this hypothesis will need further experimental work, especially after genetic accessibility of the new phylum is achieved.

**Fig. 6.**
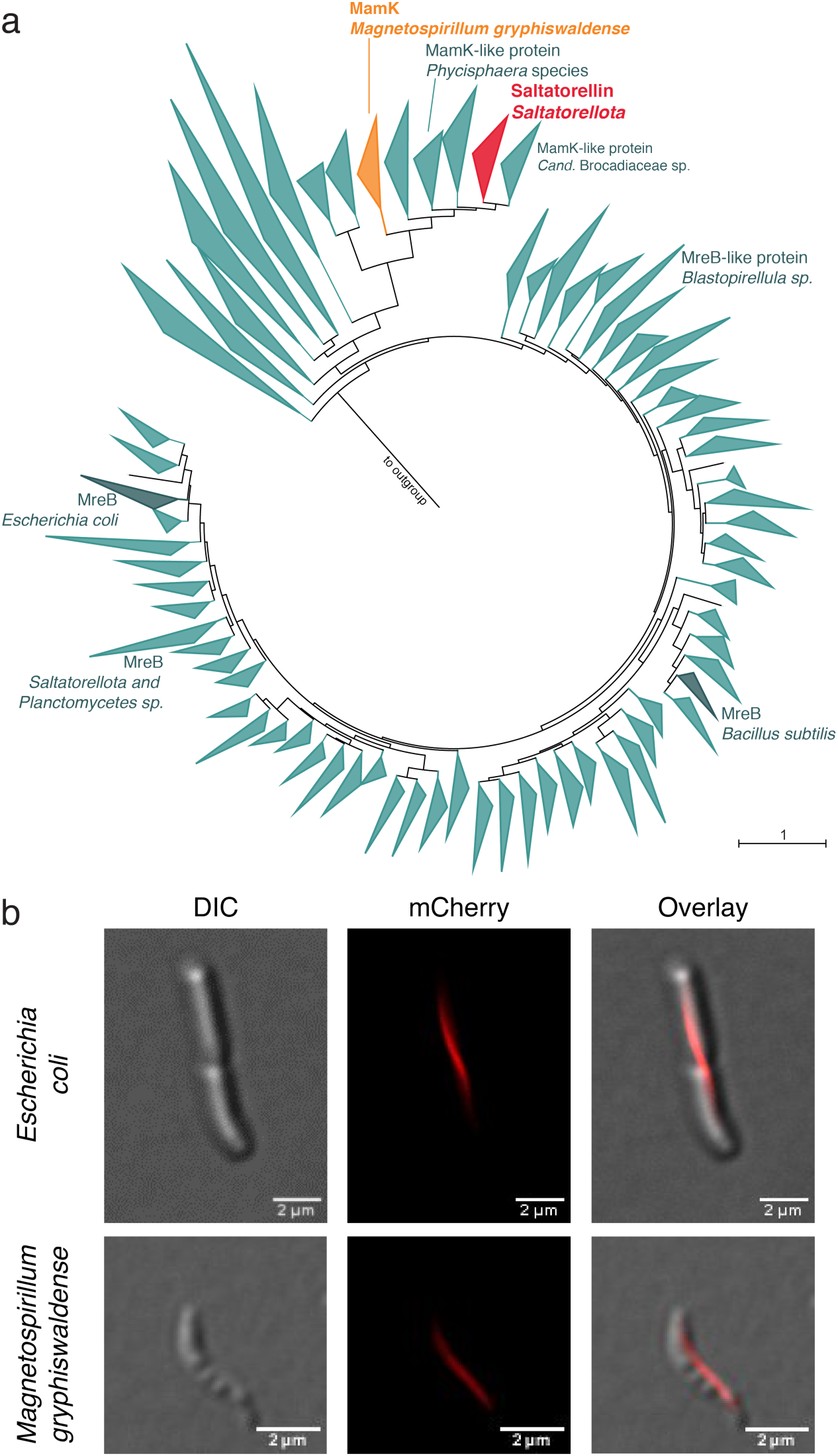
Actin homologs in *Saltatorellota*. **a)** Analysis of 14299 bacterial actin homologs, with a depicted subset of actin homologs closely related to MreB. While MreB is encoded by different species throughout the bacteria domain as expected, a second actin homolog of *Saltatorellota* branches close to MamK usually found in magnetotactic bacteria. **b)** Localization of the *Saltatorellota* MamK-like protein. Fluorescence micrographs of *E. coli* WM3064 (upper panel) and *M. gryphiswaldense* (lower panel) expressing the MamK-like from *Saltatorellus ferox* Poly30^T^ using the plasmid pBBR1-P_*tet*_-mCherry::*mamK*-like_Poly30_. In both organisms, the labelled protein localized along a filamentous structure.

### Phylogeny

From our recent publication [47] that initially mentions the three novel isolates and their genomes it does not become clear at which position within the planctomycetal phylum these novel isolates cluster as their position shifts with the method applied. To reliably determine the phylogenetic affiliation of the isolates we strived to assess them more comprehensively. At first, we appended their 16S rRNA genes to the alignment of all sequences denominated as *Planctomycetes*, *Verrucomicrobia*, *Chlamydiae*, *Lentisphaerae*, *Kiritimatiellaeota* and *Omnitrophicaeota* provided in the SILVA database release SSU Ref NR 99 [64]. Upon maximum likelihood tree formation, it became apparent that strains Poly30^T^, Pla133^T^ and Pla163^T^ cluster within the uncultured SILVA taxonomy group OM190 [32, 34] (Fig. S7), classified as subclade of the *Planctomycetes*. Although the clade appears to be monophyletic, we were not able to conclusively determine its placement within the phylum *Planctomycetes*. We found the group to either cluster together with *Candidatus* Brocadiaceae or to branch deeper or less deep. An even more random branching of the members of clade OM190 was observed when the same approach was repeated with 23S rRNA genes. A more robust phylogeny for the PVC superphylum was gained using the alternative phylogenetic marker protein RpoB (Fig. S8 and S9). With this approach, a recurring clustering of the superphylum could be achieved. A monophyletic clade comprising strains Poly30^T^, Pla133^T^ and Pla163^T^ and several metagenome-assembled genomes (MAGs) can be found in this tree. For a genome-based evaluation of the PVC superphylum, we relied on a phylogenetic assessment calculated from 120 ubiquitous single-copy proteins with the method used to build the GTDB taxonomy [65]. This again established a robust phylogenetic tree of the superphylum which contains a monophyletic clade comprising our three novel isolates as well as the GTDB taxa of uncultured MAGs UBA1135, UBA8108 and GCA-002687715 (Fig. 7a). The clusters formed by this tree are very similar to those proposed by the RpoB-based phylogenetic pattern.

**Fig. 7.**
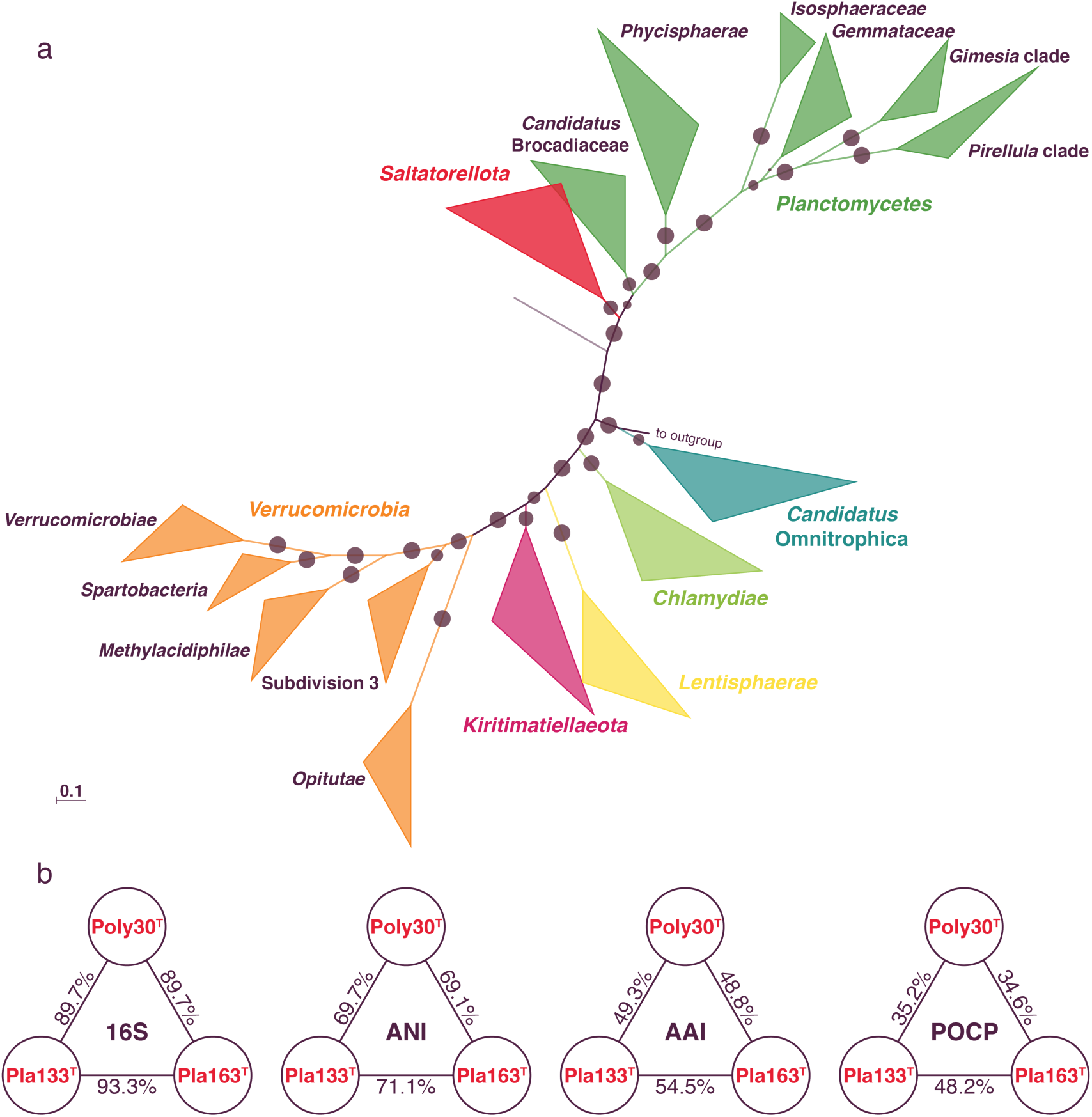
Phylogeny of the PVC superphylum and taxonomic rank determination of *Saltatorellota* strains Poly30^T^, Pla133^T^ and Pla163^T^. a) Maximum likelihood phylogeny of a genome-based alignment from 120 concatenated protein marker genes of 1675 genomes. The different colors indicate the different phyla of the PVC superphylum. Subclades of the phyla *Verrucomicrobia* and *Planctomycetes* are indicated by equal coloring. The circles indicate reliability estimators (between 0.121 −1) based on Shimodaira-Hasegawa testing (see Material and Methods). b) (From left to right) 16S rRNA sequence identity values; whole genome-based *average nucleotide identity* (ANI) values; genome-encoded protein-based *average amino acid identity* (AAI) values; genome-encoded protein-based *percentage of conserved proteins* (POCP). 16S rRNA sequence identity values >86.5% and <94.5% indicate that the tested strains belong to the same taxonomic family. ANI values <95% suggest that the compared strains belong to at least different species and AAI values <60% and POCP values <50% suggest different genera.

Until today, there are no clear standards for the designation of higher bacterial taxa, but universal thresholds based on 16S rRNA sequence identity [66] have been introduced and were used within the PVC superphylum before [67]. When applying the proposed 75% minimal identity threshold to a planctomycetal identity matrix [47], we found that strains

Poly30^T^, Pla133^T^ and Pla163^T^ do not seem to belong to the phylum *Planctomycetes* (Table S5). They might rather be the first isolates of SILVA taxon OM190 that - by the given threshold - would represent a novel, independent phylum (Table S6). A more comprehensive, whole-genome approach is provided by the relative evolutionary divergence (RED) value that is based on concatenated protein phylogeny and is the foundation of the GTDB taxonomy [65]. In this context, a RED value of 0.326±0.1 is proposed to indicate the rank of phylum. In their newest GTDB release (R04-RS89), the authors suggest the phylum Planctomycetota with a RED value of 0.227 which would also contain our isolates. However, the RED value for the determined robust cluster comprising strains Poly30^T^, Pla133^T^ and Pla163^T^ as well as UBA1135, UBA8108 and GCA-002687715 (see above) is 0.307 (Fig. S10). In contrast to the Planctomycetota value, this result is very distinctively within the given phylum threshold, thereby maintaining parity with other bacterial taxa, and hence allows to split this cluster of to a phylum in its own right. Based on the 16S rRNA gene and whole-genome analyses, and inline with the unusual traits observed in our three isolates, we suggest the novel phylum *Saltatorellota* which comprises the SILVA taxonomy phylum OM190 and the GTDB taxa UBA1135, UBA8108 and GCA-002687715 (Fig. S11). Interestingly, our calculations of 16S rRNA sequence identity and RED values imply that *Candidatus* Brocadiaceae as well as the class *Phycisphaerae* could also be transferred to novel phyla separate from the *Planctomycetes* (Table S7). However, this is beyond the scope of the present study.

To distinguish between lower taxa, 16S rRNA sequence identities can be used alongside average nucleotide identities (ANI), average amino acid identities (AAI) and the percentage of conserved proteins (POCP). For strains Poly30^T^, Pla133^T^ and Pla163^T^, the determined values (Fig. 7b) imply that all three isolates belong to different species as well as different genera. This rating is seconded by 16S rRNA sequence identities, which additionally allow the conclusion that all strains belong to one taxonomic family.

On the basis of this phylogenetic analysis, we suggest that strains Poly30^T^, Pla133^T^ and Pla163^T^ represent novel species of novel genera for which we propose the names *Saltatorellus ferox* sp. nov, gen. nov., *Engelhardtia mirabilis* sp. nov, gen. nov and *Rohdeia mirabilis* sp. nov, gen. nov. With *Saltatorellus* being the type genus, all three genera belong to the family *Saltatorellaceae* within the new phylum *Saltatorellota* [68]. The *Saltatorellota* phylum comprises the SILVA taxonomy phylum OM190 and the GTDB taxa UBA1135, UBA8108 and GCA-002687715.

### Description of *Saltatorellus* gen. nov

*Saltatorellus* (Sal.ta.to.rel’lus. N.L. masc. n., *Saltatorellus* dim. of L. saltator a dancer; a bacterium that occurs to be dancing). The type species of the genus is *Saltatorellus ferox*.

### Description of *Saltatorellus ferox* sp. nov

*Saltatorellus ferox* (fe’rox. L. masc. adj. ferox boisterous, wild; corresponding to the wild movements of the cells). *Saltatorellus ferox* cells show amoeba-like locomotion and shapeshifting. If cells form large aggregates the formation of trunk-like structures was observed. Cell size is very variable from 0.8-3.5 µm with an average cell size of 1-1.3 µm (Fig. S12). The GC content is 67%. *Saltatorellus ferox* grows in a range of artificial seawater (ASW) between 10-100% with optimal growth between 25%-75%. NaCl of 0% and 1% was tolerated while at 2.5% and higher concentration no growth was observed. *Saltatorellus ferox* grows at temperatures between 15-33°C with optimal growth at 27 - 30°C. Cells showed similar growth at pH 7 and pH 7.5 while below and above these pH values no growth was observed.

The type species is strain Poly30^T^ (= DSM 103386) isolated from polyethylene particles incubated near the shore of Heiligendamm, Germany in the Baltic Sea (54.146 N 11.843 E). The GenBank accession numbers are MK559991 for the 16S rRNA gene sequence and CP036434 for the genome sequence.

### Description of *Rohdeia* gen. nov

*Rohdeia* (Roh.de’i.a. N.L. fem. n. Rohdeia, a bacterium named in honor of the German microbiologist and electron microscopist Manfred Rohde, who contributed significantly to the visualization of adhesion and invasion mechanisms of pathogenic bacteria, to the morphological characterization of bacteriophages and newly isolated bacteria, to the advancement of various TEM and SEM methods as well as to the cell biological characterization of the phylum *Planctomycetes*). *Rohdeia mirabilis* is the type species of this genus.

### Description of *Rohdeia mirabilis* sp. nov

*Rohdeia mirabilis* (mi.ra’bi.lis. L. fem. adj. mirabilis marvelous, wonderful; corresponding to special attributes of the strain). *Rohdeia mirabilis* cells grow in huge aggregate of millimeter size. Its GC content is 70.5%. *Rohdeia mirabilis* growth in a temperature range of 20-30 °C with optimal growth at 27 °C.

The type species is strain Pla163^T^ isolated from biofilms on incubated wood or plastic particles in the Baltic Sea near an inflowing river estuary at the coastline of Rostock (Hohe Düne), Germany (54.183 N 12.095 E). The GenBank accession numbers are MK559986 for the 16S rRNA gene sequence and CP036290 for the genome sequence.

### Description of *Engelhardtia* gen. nov

*Engelhardtia* (En.gel.hard’ti.a. N.L. fem. n. Engelhardtia, named in honor of the German structural microbiologist Harald Engelhardt, who contributed significantly to the structural understanding of the archaeal and bacterial cell envelope, to the advancement of cryo-electron tomography as well as to the cell biological characterization of the phylum *Planctomycetes*). *Engelhardtia mirabilis* is the type species of the genus.

### Description of *Engelhardtia mirabilis* sp. nov

*Engelhardtia mirabilis* (mi.ra’bi.lis. L. fem. adj. mirabilis marvelous, wonderful; corresponding to special attributes of the strain). *Engelhardtia mirabilis* cells are in average 0.8-1.3 µm in diameter, but sizes can vary between 0.5-2.0 µm (Fig. S12). They possess multiple interconnected cytoplasmic invaginations. Its GC-content is 70.3%. *Engelhardtia mirabilis* growth in a temperature range of 20-33 °C with optimal growth at 30 °C. It tolerates a pH range of 6.5 - 7.5 with optimal growth at pH 7.

The type species is strain Pla133^T^ (= DSM 103766) isolated from polystyrene particles incubated at the Warnow river estuary in Rostock, Germany (54.106 N 12.096 E). The GenBank accession numbers are MK559985 for the 16S rRNA gene sequence and CP036287 and CP036288 for the genome sequences.

### Description of *Saltatorellaceae* fam. nov

*Saltatorellaceae* (*Sal.ta.to.rel.la.*ce’ae. N.L. masc. dim. n. *Saltatorellus*, type genus of the family; suff. *-aceae*, ending to denote a family; N.L. fem. pl. n. *Saltatorellaceae*, the *Saltatorellus* family). The type genus of the family is *Saltatorellus*.

### Description of *Saltatorellales* ord. nov

*Saltatorellales* (*Sal.ta.to.rel.la’les*. N.L. masc. dim. n. *Saltatorellus*, type genus of the order; suff. *-ales*, ending to denote an order; N.L. fem. pl. n. *Saltatorellales*, the *Saltatorellus* order). The type genus of the order is *Saltatorellus*.

### Description of *Saltatorellae* class nov

*Saltatorellae* (*Sal.ta.to.rel’lae*. N.L. masc. dim. n. *Saltatorellus*, type genus of the type order of the class; N.L. fem. pl. n. *Saltatorellae*, the class of the order *Saltatorellales*). The type order of the class is *Saltatorellales*.

### Description of *Saltatorellota* phyl. nov

*Saltatorellota* (*Sal.ta.to.rell’ota.* N.L. fem. pl. n. *Saltatorellae*, type class of the phylum; suff. *-ota*, ending to denote a phylum; N.L. neut. pl. n. *Saltatorellota*, the phylum of the class *Saltatorellae*). Based on a phylogenetic assessment, calculated from 120 ubiquitous single-copy proteins, the proposed phylum *Saltatorellota* forms a stable monophyletic linage separated from the phylum *Planctomycetes*. Similar results were obtained for a RpoB-based phylogeny. While a 16S rRNA gene based phylogeny showed no reliable clustering, the 16S rRNA gene sequence identity between members of the proposed phylum *Saltatorellota* and the phylum *Planctomycetes* is below the recommended threshold of 75%, supporting separate phyla [66]. The proposed phylum *Saltatorellota* includes the uncultured GTDB taxa UBA1135, UBA8108 and GCA-002687715 and SILVA taxon OM190. The type class of the phylum is *Saltatorellae*.

## Conclusion

Some conclusions and evolutionary implications that one might think of when looking at the phylum *Saltatorellota* might parallel the historical view on the phylum *Planctomycetes [2]* that were once considered the ‘missing link’ between eukaryotes and prokaryotes/bacteria [2] - a claim we do not want to make in this manuscript. This is why we concentrated on the description of the novel phylum with a focus on controls such as determining the natural growth conditions. The work was performed in six independent research centers and all critical time-lapse analyses were done in parallel in two different laboratories employing two different experimental setups. Thus, we are confident in the significance of our observations. Nevertheless, given the history of the PVC superphylum -from the *Chlamydia* anomaly to the paradigm shift in planctomycetal cell biology – we will not speculate about potential evolutionary implications of our findings, but stress the need for further analysis requiring the development of genetic tools to reveal the molecular background of the observed traits.

## Materials and Methods

### Isolation, cultivation and physiological characterization of *Saltatorellota* strains

For the isolation of *E. mirabilis* Pla133^T^ and *R. mirabilis* Pla163^T^, microplastic and wood particles collected on 04 September 2014 were incubated for 16 days in an inflowing river estuary and at the Baltic Sea coast as previously described [57]. *S. ferox* Poly30^T^ was isolated from plastic particles collected on 08 October 2015 (incubation time two weeks) from the Baltic Sea at Heiligendamm pier, Germany. Details concerning isolation strategy and media were previously described [47].

All growth tests were measured in triplicates. Growth at pH 5-9 in 0.5 intervals was tested using 100 mM MES, HEPES, HEPPS or CHES buffer, corresponding to the individual buffer range. Growth was tested at 10, 15, 20, 22, 24, 27, 30, 33, 36 and 40 °C to identify the optimal growth temperature. The optimal salinity for *S. ferox* Poly30^T^ was tested by inoculation of artificial seawater (ASW) at concentrations of 5%, 10% and 25-300% in 25% intervals to test if the strains are growing under hypersaline conditions. 50% ASW corresponds to 250 ml/L concentrated artificial sea water (46.94 g/L NaCl, 7.84 g/L Na_2_SO_4_, 21.28 g/L MgCl_2_ x 6 H_2_O, 2.86 g/L CaCl_2_ x 2 H_2_O, 0.384 g/L NaHCO_3_, 1.384 g/L KCl, 0.192 g/L KBr, 0.052 g/L H_3_BO_3_, 0.08 g/L SrCl_2_ x 6 H_2_O and 0.006 g/L NaF). NaCl tolerance for *S. ferox* Poly30^T^ was tested between 0-10% at 0%, 1%, 2.5%, 3,5%, 4% and in 1% intervals at higher concentrations. Therefore, medium with 50% ASW (standard cultivation conditions) was used and only the NaCl concentration was varied.

### Baltic Sea environmental sample collection along a salinity gradient

Near surface water samples (5m depth) for determination of *in situ* relative abundances of *Saltatorellota* were taken in Aug/Sep 2015 onboard the R/V Poseidon (cruise POS488) along the southern and eastern coastlines of the Baltic Sea, from Rostock, Germany to Helsinki, Finland (Fig. 5). Samples were collected in 5 L Free-Flow bottles (Hydro-Bios) mounted on a rosette equipped with a Conductivity-Temperature-Depth-probe (Sea-Bird SBE 9). Water from 5 - 6 Free-Flow bottles was mixed and filtered in technical triplicates (500 mL each) through a 3 µm pore-size cellulose nitrate filter (particle-attached fraction) and a 0.22 µm pore cellulose nitrate filter (both GE Whatman) to capture the free-living bacterial fraction. Filters were snap-frozen in liquid nitrogen and stored at −80 °C until further processing.

### DNA extraction, 16S rRNA-gene amplicon sequencing and sequence processing

DNA from the collected filter was extracted using the DNeasy PowerSoil Kit (Qiagen) according to the manufacturer’s instructions, except that DNA was eluted twice from the spin column using the same 50 µL PCR-grade water. Amplification of DNA was conducted via polymerase chain reaction (PCR) using bacterial primers covering the V4-region of the 16S rRNA-gene (position 515F to 806R in *E. coli*) with the forward sequence 5’ GTGCCAGCMGCCGCGGTAA 3’ and reverse sequence 5’ GGACTACHVGGGTW TCTAAT 3’ [69] PCR conditions were as followed; initial denaturation at 98 °C for 2 min, followed by an additional denaturation step at 98 °C for 15 s, annealing at 65 °C for 15 s and elongation for 30 s at 68 °C. Steps 2 – 4 were repeated 9 times, with elongation temperature < 1 °C per cycle (linear amplification). After 9 cycles of linear amplification, this was followed by a denaturation step at 98 °C for 15 s, annealing at 55 °C for 15 s and elongation at 68 °C for 30 s for 24 cycles and a final elongation step at 70 °C for 5 min [70]. Library preparation and sequencing on an Illumina MiSeq machine were carried out according to the ‘Illumina 16S Metagenomic Sequencing Library Preparation Guide’ at the Institute of Medical Microbiology, Virology and Hygiene, University Medical Centre Rostock, Germany. Raw sequence reads are available from the NCBI Sequence Read Archive under the accession number PRJNA506548 (condition = *in situ*).

### Sequence processing and filtering for sequences of cultivated *Saltatorellota* strains

Obtained sequences were processed using the mothur-pipeline v. 1.39.5 [71]. Paired end reads were assembled, and quality filtered (permitted length = 250 – 275 bp, max. number of ambiguous bases per sequence = 0, max. number of homopolymers per sequence = 8). Sequences were classified using the Wang algorithm [72] (required bootstrap value of ≥ 85%) and the SILVA SSURef release 132 as reference database [73], into which the sequences of the complete 16S rRNA gene of the strains Poly30_41280, Pla163_29910, and Pla133_39970 (GenBank acc. no. MK559985, MK559986 and MK559991) had been included. Sequences classified as *Eukaryota*, *Archaea*, Chloroplasts, Mitochondria, and as unknown were excluded from the dataset. *Saltatorellota* sequences were extracted using the ‘get.lineages’ command in mothur and OTUs picked from the sequences at a 97% threshold. Finally, OTUs were classified, relative abundances calculated based on the library size of bacterial reads per sample, and mean abundances per station calculated for all *Saltatorellota* OTUs together. Mean relative abundances of particle-attached (> 3 µm) *Saltatorellota* spp. at the different cruise stations were visualized using Ocean Data View (Schlitzer, R., Ocean Data View, odv.awi.de, 2018). Salinity values for the coastal zones were extrapolated from the measured *in situ* data within Ocean Data View using Data Interpolating Variational Analysis (DIVA) with default parameters. Salinity and relative abundance maps were merged using the graphical software Gimp v. 2.10.8.

### Phylogenetic analysis of the PVC phylum and *Saltatorellota*

To resolve the placement of strains Pla163^T^, Pla133^T^ and Poly30^T^ within the phylogeny of the PVC superphylum by means of 16S and 23S rRNA-based analysis, alignments for this clade were obtained from the non-redundant SILVA SSU Ref NR 99 database, release 132 or the SILVA LSU Ref, release 132, respectively [73]. 16S and 23S rRNA sequences from the novel strains (GenBank acc. no. MK559985, MK559986 and MK559991) were aligned with SINA [74] and appended. Ten sequences derived from different *Proteobacteria*, *Firmicutes* and *Gemmatimonadetes* were used as outgroup. Different trees were determined by using different cut-off positions when cropping the alignment and by using a 50% identity filter. Maximum likelihood inference was calculated employing FastTree v2.1.10 [75] with the GTRGAMMA substitution model invoked. A subset alignment containing only sequences of SILVA taxonomy group OM190 and the three novel strains was created (positions 5295-41022). The maximum likelihood inference for these trees was derived by using RAxML [76] (1,000 bootstraps, model of nucleotide substitution: GTRGAMMAI). Ten sequences derived from different *Proteobacteria*, *Firmicutes* and *Gemmatimonadetes* and all planctomycetal 16S rRNA sequences published recently by Wiegand *et al.* [47] were used as outgroup. An identity matrix of this alignment was created with ARB [77].

For the analyses based on whole genomes or single genes genome data were obtained from NCBI GenBank based on the NCBI Taxonomy database. All available genomes classified as members of the PVC phylum were downloaded on 7 January 2019. Completeness and contamination of all genomes was determined using CheckM v1.0.13 [78]. Only genomes with a completeness >50%, a contamination <10%, and a strain heterogeneity 0 were included in the following analyses. Additionally, all planctomycetal genomes (including the *Saltatorellota* genomes wit GenBank acc. no. CP036287+CP036288, CP036290 and CP036434) sequenced by Wiegand *et al. [47]* were included and ten genomes from *Proteobacteria*, *Firmicutes* and *Gemmatimonadetes* served as outgroup.

Phylogenetic placements based on the marker protein RpoB were computed based on the aforementioned genomes from which RpoB protein sequences were extracted by using hmmer [79] together with the TIGR02013 alignment. The results were aligned with MUSCLE v.3.8.31 [80], the ends of the alignment were curated, and the phylogeny was inferred by maximum likelihood implemented in FastTree v2.1.10 [75] (WAG substitution model and gamma distributed rate heterogeneity enabled). An additional phylogeny was inferred from RpoB sequences only belonging to the three novel strains, one member of each potential planctomycetal genus [47] and three *Verrucomicrobia* as outgroup. For this smaller subset the alignment was made with Clustal Omega [81] and the phylogenetic tree was calculated with a) RAxML [76] and the WAG substitution model and gamma distributed rate heterogeneity enabled (1,000 bootstrapping resamples) and b) Bayesian interference as implemented in Mr Bayes [82] (rate variation model: invgamma, amino acid model: fixed(wag), single chain per analysis, number of generations: 300,000).

A genome-based alignment relying on marker gene sets was gained by employing GTDB-Tk v0.2.2 [65] (https://github.com/Ecogenomics/GtdbTk/) with the ‘identify’ and ‘align’ option. The phylogeny was inferred with the maximum likelihood model implemented in FastTree v2.1.10 [75] (gamma distributed rate heterogeneity and Shimodaira-Hasegawa support values enabled) and using different amino acid substitution models to test for robustness (WAG, JTT and LG). For Fig. 7, the tree calculated with WAG substitution level was used. Each analysis was performed for all included sequences and for subsets, each with one taxon removed. RED values, that have also been introduced by Parks *et al.* [65], were determined by employing PhyloRank v0.0.37 (https://github.com/dparks1134/PhyloRank) on the above used alignment and the taxonomy file gained from GTDB-Tk. Editing the taxonomy file allowed to determine custom RED values for newly defined taxa.

A more custom approach, as used by Bartling *et al.* [83] and Vollmers *et al.* [33], to genome-based phylogeny that does not rely on marker genes was used for members of the above defined taxon comprising the novel strains Pla163^T^, Pla133^T^ and Poly30^T^. In this analysis, three planctomycetal strains served as outgroup. The unique single-copy core genome of all genomes was determined with proteinortho5 [84] using the following parameters "-cov=55-e=1e-5-identity=25" and tolerating the absence of single-copy genes in up to two comparison genomes to account for partial draft genomes. The protein sequences of the resulting orthologous groups were aligned using MUSCLE v.3.8.31 [80]. After clipping partially aligned C- and N-terminal trailing regions, poorly aligned internal alignment regions were subsequently filtered using Gblocks [85]. To reduce influence from potential horizontal gene transfer events, each orthologous group was then preliminarily clustered using FastTree v2.1.10 [75], and the resulting tree topologies were compared with respect to the observed summed phylogenetic distances. Markers which produced tree topologies with summed phylogenetic distances larger than the mean for all orthologous groups plus two standard deviations were dismissed. The final selection of seven marker alignments with a combined length of 1392 conserved amino acid residues was concatenated and clustered using the maximum likelihood method implemented by RAxML [76] with the "rapid bootstrap" method and 100 bootstrap replicates. To generate similarity matrices, the average amino acid (AAI) was gained with the aai.rb script of the enveomics collection [86], the average nucleotide identity (ANI) was determined by OrthoAni [87] and the percentage of conserved proteins (POCP) was calculated as described by Qin *et al.* [88]. All trees were collapsed and formatted with iTOL v4 [89].

### Presence of genes involved in swimming, twitching, gliding and sliding motility

The targets for the analysis were chosen from literature [58, 63, 90-96]. (Pfam [97] and TIGRFAM [98] identifiers were gathered for all genes from the UniProt database [99] (Table S2). Profile Hidden Markov Models (HMM) were computed from Pfam rp55 and TIGRFAM seed alignments with hmmbuild [79]. The profile HMMs were then used to search genomes of strains Pla133^T^, Pla163^T^ and Poly30^T^ with hmmsearch [79]. Gene clusters with potential hits were gathered and locally re-evaluated with InterProScan [100]. The results were manually curated (Table S2) and transferred into Fig. S4 and S5 and Table S3.

### Analysis of motor protein-homologs

The targets for the analysis were the eukaryotic motor proteins myosin (PF00063), kinesin (PF00225) and dynamin (PF00350). Profile Hidden Markov Models (HMM) were computed from the Pfam rp55 seed alignments with hmmbuild [79]. The profile HMMs were then used to search genomes of strains Pla133^T^, Pla163^T^ and Poly30^T^ with hmmsearch [79].

### Phylogeny of bacterial actin homologs

For determining the phylogeny of bacterial actin homologs such as MreB, HMM profiles built with the sequences found by Derman *et al.* [101] was used for a hmmsearch [79] analysis against all sequences in the BLAST nr database (June 2018). Additionally, a BLASTp analysis of all sequences was conducted against the BLAST nr database. The results were merged, deduplicated and chaperone DnaK (*E. coli*) and ribulokinase AraB (*E. coli*) were blasted (BLASTp) against them. This was done as both proteins show similarities to the actin homologs but do not form filaments themselves (see for example CDD-Acc. cd00012 [102]). All hits that had better hits against DnaK or AraB were removed. The remaining sequences were used as query sequences in a DIAMOND [103] analysis against the BLAST nr database. Again, the results were deduplicated and DnaK and AraB hits were removed. The process was repeated for another two cycles.

All sequences <100 aa or >800 aa were removed from the resulting amino acid sequence file (48443 entries). Subsequently, the sequences were clustered with USEARCH [104] and an identity threshold of 50% (‘-sort length’ option enabled). The representative sequences of each cluster were aligned with MAFFT [105] (with the ‘--retree 2’ and ‘--maxiterate 0’ options enabled) together with reference sequences: the amino acid sequences from Derman *et al.* [101], all planctomycetal actin homolog hits (see Wiegand *et al.* [47]), DnaK, AraB and DnaA as outgroup. A phylogenetic tree was built with FastTree2 [75]. One subset of the tree was chosen and the sequences as well as all sequences from their respective USEARCH clusters were extracted. With this gene set, a second iteration was started as described above (with an USERACH identity threshold of 75% and ‘-sort’ option disabled). A third (with an USERACH identity threshold of 85% and ‘-sort’ option disabled) and a fourth (with an USERACH identity threshold of 85% and ‘-sort length’ option enabled, additional amino acid sequences of Mollicutes *mreB*s as reference) refinement followed.

### GC-MS analysis of diaminopimelic acid from *Saltatorellota* cells

Detection was performed as previously described [38]. In brief, cells were harvested from 6 - 20 mL cultures by centrifugation, lyophilized, hydrolyzed (200 µl 4 N HCl, 100 °C, 16 h) and dried in a vacuum desiccator. Amino acids derivatized to N-heptafluorobutyryl isobutylesters [106] were resolved in ethyl acetate and analyzed by GC/MS (Singlequad 320, Varian; electron impact ionization, scan range 60 to 800 m/z). The derivatized diaminopimelic acid was detected in Extracted Ion Chromatograms (EIC) using the characteristic fragment ion set 380, 324, 306 and 278 m/z at a retention time of 23.5 – 23.55 min.

### TEM and SEM of *Saltatorellota* cells and sacculi

For field emission scanning electron microscopy (FESEM) *Saltatorellota* cells were fixed in 1% formaldehyde in HEPES buffer (3 mM HEPES, 0.3 mM CaCl_2_, 0.3 mM MgCl_2_, 2.7 mM sucrose, pH 6.9) for 1 h on ice and washed one time with HEPES buffer. Cover slips with a diameter of 12 mm were coated with a poly–L–lysine solution (Sigma–Aldrich) for 10 min, washed in distilled water and air–dried. 50 µl of the fixed bacteria solution was placed on a cover slip and allowed to settle for 10 min. Cover slips were then fixed in 1% glutaraldehyde in TE buffer (20 mM TRIS, 1 mM EDTA, pH 6,9) for 5 min at room temperature and subsequently washed twice with TE–buffer before dehydrating in a graded series of acetone (10, 30, 50, 70, 90, 100%) on ice for 10 min at each concentration. Samples from the 100% acetone step were brought to room temperature before placing them in fresh 100% acetone. Samples were then subjected to critical-point drying with liquid CO_2_ (CPD 300, Leica). Dried samples were covered with a gold/palladium (80/20) film by sputter coating (SCD 500, Bal– Tec) before examination in a field emission scanning electron microscope (Zeiss Merlin) using the Everhart Thornley HESE2–detector and the inlens SE–detector in a 25:75 ratio at an acceleration voltage of 5 kV.

Cell pellets were resuspended in 10 mL 10 % SDS (w/v) and boiled for three hours. Sacculi were harvested (30 min, 139 699 x g, 4 °C; SW 60 Ti rotor, Beckman Coulter), washed 4 times in 3 mL water and resuspended in 1 mL water supplemented with 0.02% (w/v) sodium azide. TEM micrographs of murein sacculi were taken after negatively staining with 2% aqueous uranyl acetate, employing a Zeiss transmission electron microscope EM 910 at an acceleration voltage of 80 kV at calibrated magnifications as previously described [107].

### Cryo-electron tomography of *Engelhardtia mirabilis* Pla133^T^

Cells of strain Pla133^T^ were mixed with the same volume of BSA-stabilized 15 nm colloidal gold solution (Aurion) and placed on holey carbon-coated 200 mesh copper grids (R2/1, Quantifoil, Jena, Germany) immediately before thin-film vitrification by plunge-freezing in liquid propane (63%)/ethane (37%) [108]. Typically, grids with frozen-hydrated samples were mounted in Autogrids [109]. Tomographic tilt series were recorded under low dose conditions (total dose typically <150 e/Å^2^) on a Titan Krios (FEI, Eindhoven, The Netherlands) equipped with a Bioquantum post-column energy filter with a K2 Summit direct electron detector (Gatan, Pleasanton, CA, USA). At each tilt, dose fractionation mode was employed, and 8 frames of each projection were sampled, which were then aligned to compensate for beam-induced object drift using MotionCor2 [110]. Typically, tilt series were recorded at a nominal defocus of −5 or −6 µm, and a primary magnification of 33000 x (corresponding to pixel sizes on the object level of 0.43 nm and covered an angular range of ± 60° in increments of 2°. IMOD [111] was used for 3D reconstruction.

### Time-lapse Microscopy of *Saltatorellota* cells and cell aggregates

Given the very unusual observations we made, time-lapse experiments were performed in two independent laboratories with distinct experimental settings and from two independent investigators to double check all results.

In Nijmegen, 2 µL of culture were added to the glass bottom of a Nunc Glass Base Dish. Cells were immobilized using a 1% agarose cushion consisting of M3H NAG ASW no SL10 medium and agarose. The sample was sealed using Vaseline. For imaging, environmental control was set to 28 °C. Pictures were taken every 5 minutes for a total of 24 h with a Leica DMi8 S inverse microscope using a HC PL APO 100x/1.40 OIL objective.

In Braunschweig, time-lapse microscopy was performed as described before [112]. Briefly, 2 µl of Poly30^T^ and Pla133^T^ cells of an exponentially growing culture were immobilized on a 1% agarose–pad, containing M1HNAGASW (without sl10) medium and were imaged in a MatTek Glass Bottom Microwell Dish (35 mm dish, 14 mm Microwell with No. 1.5 cover-glass P35G-1.5-14-C). Images were taken with phase–contrast illumination using a NikonTi microscope with a Nikon N Plan Apochromat λ 100x/1.45 Oil objective and the ORCA-Flash 4.0 Hamamatsu or RI2 cameras. Cell growth, locomotion, cell division and pili formation were observed every 5 min for up to 48 h at 28°C. Micrographs were subsequently aligned and analyzed using the NIS-Elements imaging software V 4.3 (Nikon).

### Heterologous expression and florescent microscopy of the mCherry tagged MamK-like protein from strain Poly30^T^

*E. coli* strain WM3064 (W. Metcalf, unpublished) was cultivated in lysogeny broth (LB) medium supplemented with 1 mM DL-α, Ɛ-diaminopimelic acid (DAP) at 37 °C and shaking (180 rpm). A *MamK-*deleted mutant strain of *M. gryphiswaldense* [113] was cultivated microaerobically at 28 °C in modified flask standard medium (FSM) [114]. For the selection of strains carrying recombinant plasmids, media were supplemented with 25 µg/mL kanamycin for *E. coli* WM3064 and 5 µg/mL for *M. gryphiswaldense*. 1.5% (w/v) agar was added for solid medium.

Oligonucleotides used for the construction of plasmid pBBR1-P_*tet*_-mCherry::*mamK*-like_Poly30_ were purchased from Sigma-Aldrich (Steinheim, Germany). The *MamK*-like gene from strain Poly30^T^ was amplified by standard PCR procedures with Phusion DNA polymerase (Thermo Scientific) using the primers RPA821_NheI (CTA<u>**GCTAGC**</u>ATGACCGACATC ACGACCGAC) and RPA822_BamHI (CGC<u>**GGATCC**</u>TCACTTGAGCTGCTCCCAGTAG). The digested PCR product was ligated with plasmid pBBR1-P_*tet*_-mCherry (pAP160 modified with mCherry [115]) containing inducible promoter P_*tet*_ digested with same enzymes, yielding pBBR1-P_*tet*_-mCherry::*mamK*-like_Poly30_. The sequence-verified (Macrogen Europe, Amsterdam, Netherlands) construct was introduced into chemically competent *E. coli* WM3064 following the standard procedure. *E. coli* WM3064 was used as a donor to transfer a recombinant plasmid into Δ*mamK* strain of *M. gryphiswaldense* by conjugation as described earlier [116].

For induction experiments, media were supplemented with 100 ng/mL anhydrotetracycline (Atet). Δ*mamK* of *M. gryphiswaldense* harboring pBBR1-P_*tet*_-mCherry::*mamK*-likePoly30 was grown under an undefined microoxic condition in 15 mL polypropylene tubes with sealed caps in a culture volume of 10 mL to early mid-log phase. *E. coli* WM3064 with pBBR1-P_*tet*_-mCherry::*mamK*-like_Poly30_ was grown under aerobic condition in a 50 mL Erlenmeyer flask with 10 mL culture volume. After 6 h of induction, 5 µL of cell culture was fixed on a ‘MSR-agarose pad’ [63] and covered with a coverslip. Fluorescence microscopy of immobilized cells was performed with an Olympus BX81 microscope equipped with a 100X UPLSAPO100XO objective (with a numerical aperture of 1.40) and an Orca-ER camera (Hamamatsu). Images were analyzed and prepared with ImageJ Fiji v1.50c [117].

## Acknowledgment

We thank Anja Heuer for skillful technical assistance in cultivation and Ina Schleicher for help with the electron microscopic preparations for FSEM and negative-staining. We further thank the captain and crew of the R/V Poseidon cruise POS488 and Bernd Kreikemeyer (Medical Microbiology, Virology and Hygiene, University Medical Centre Rostock, Germany) for providing sequencing opportunities. The technical assistance of Jana Bull for preparation of sequencing runs is greatly acknowledged.

## Data and materials availability

All data is available in the main text or the supplementary materials.

