## SupplementaryInformation for "The novel shapeshifting bacterial phylum *Saltatorellota*"

### Supplementary Figures

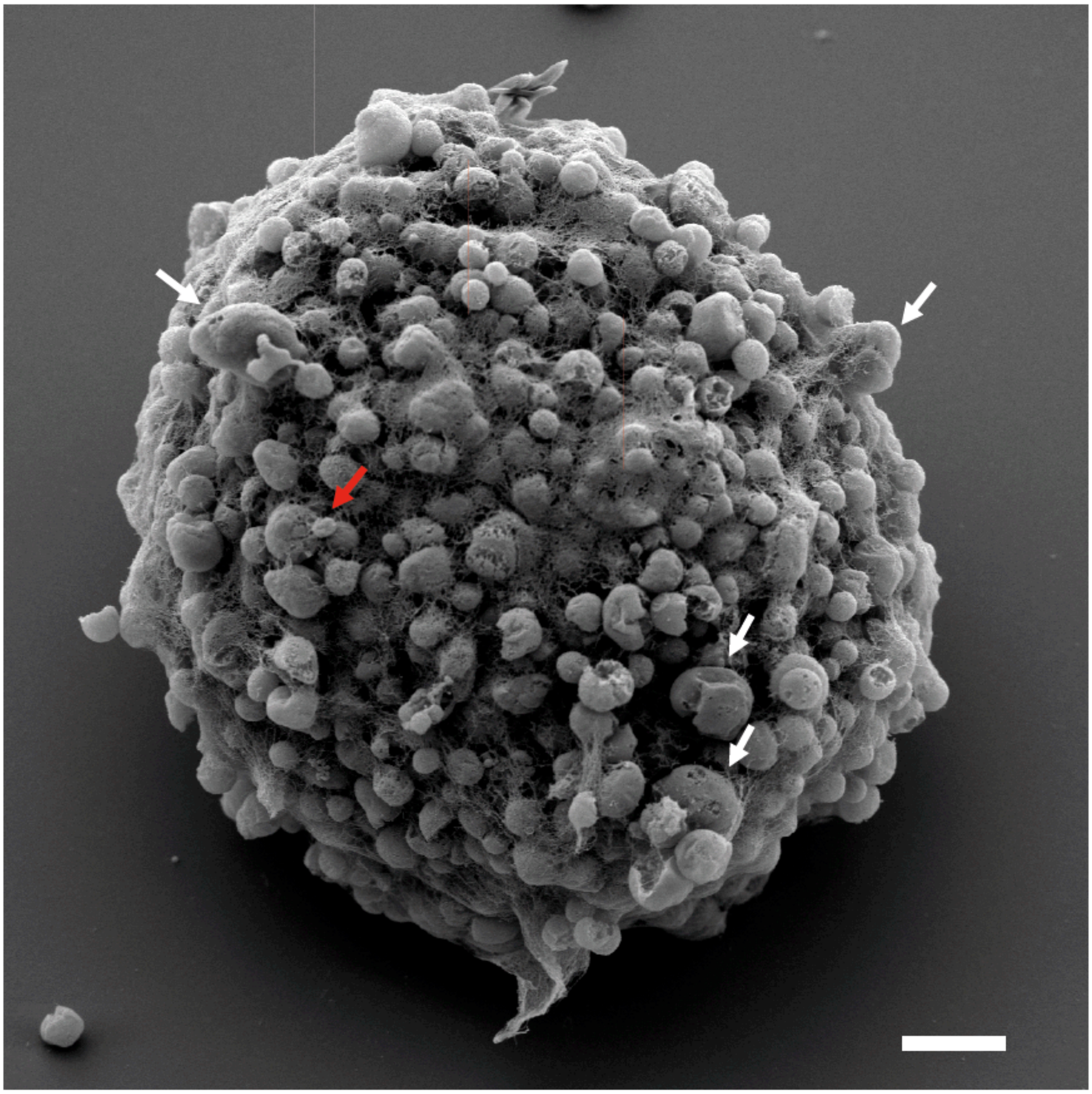

**Fig. S1. *Saltatorellota* strains tend to form aggregates.** SEM analysis revealed that aggregate forming *Saltatorellus ferox* Poly30$^T$ displays extreme polymorphism with giant amorphic- (white arrows) and small coccoid cells (red arrows). Scale bar 3 µm.

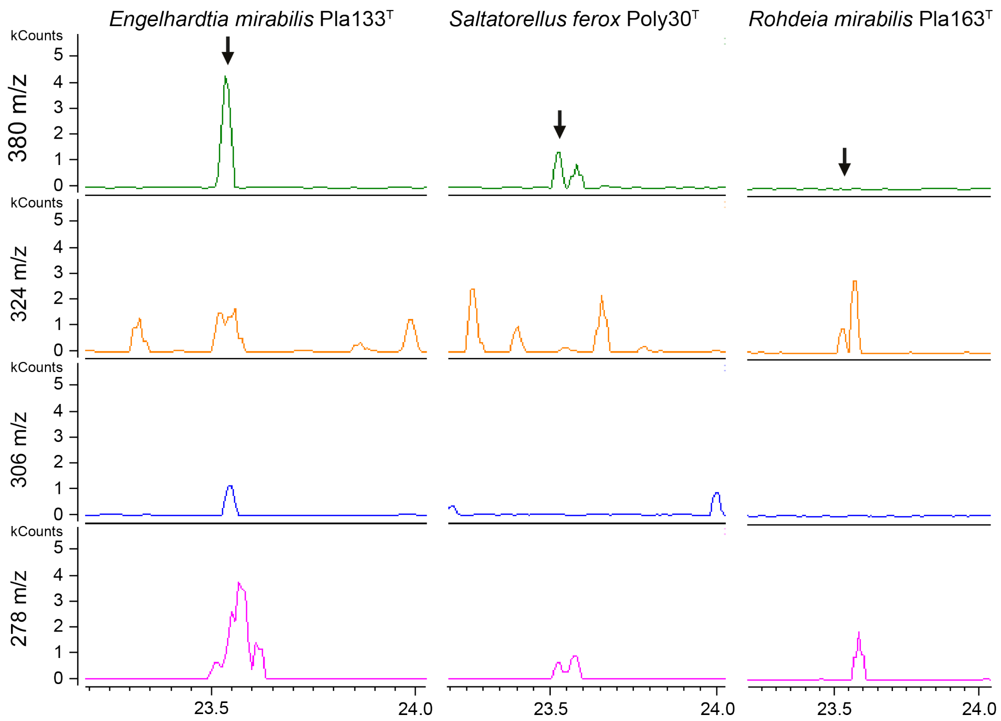

**Fig. S2. Mass spectrometric detection of diaminopimelic acid (DAP) from *Saltatorellota* cells**. Extracted ion chromatograms of DAP derivative (N-heptafluorobutyryl DAP isobutylester) from whole-cell hydrolysates of *Engelhardtia mirabilis* Pla133$^{T}$, *Saltatorellus ferox* Poly30$^{T}$ and *Rohdeia mirabilis* Pla163$^{T}$. Masses of DAP fragments (380, 324, 306, and 278 m/z) correspond to peaks at 23.5-23.55 min retention time, indicated by black arrows in the top panel.

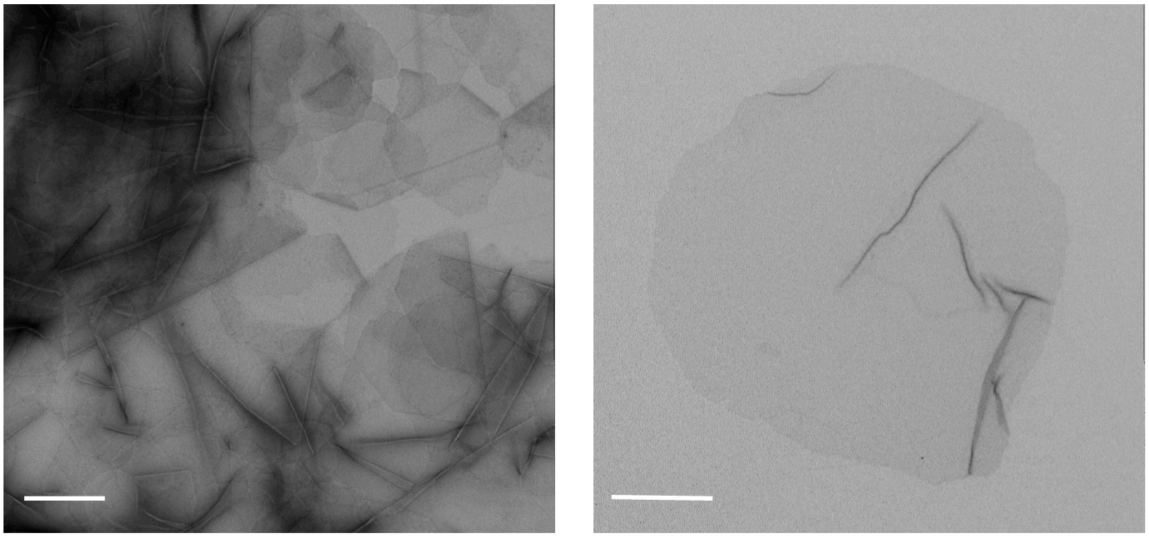

**Fig. S3. Peptidoglycan sacculi**. Peptidoglycan sacculi were obtained from *Rohdeia mirabilis* Pla163$^{T}$ after boiling cells in 10% SDS. (Left) TEM micrograph showing an accumulation of negative stained sacculi. (Right) TEM micrograph of an individual negative stained sacculus. Scale bar 200 nm.

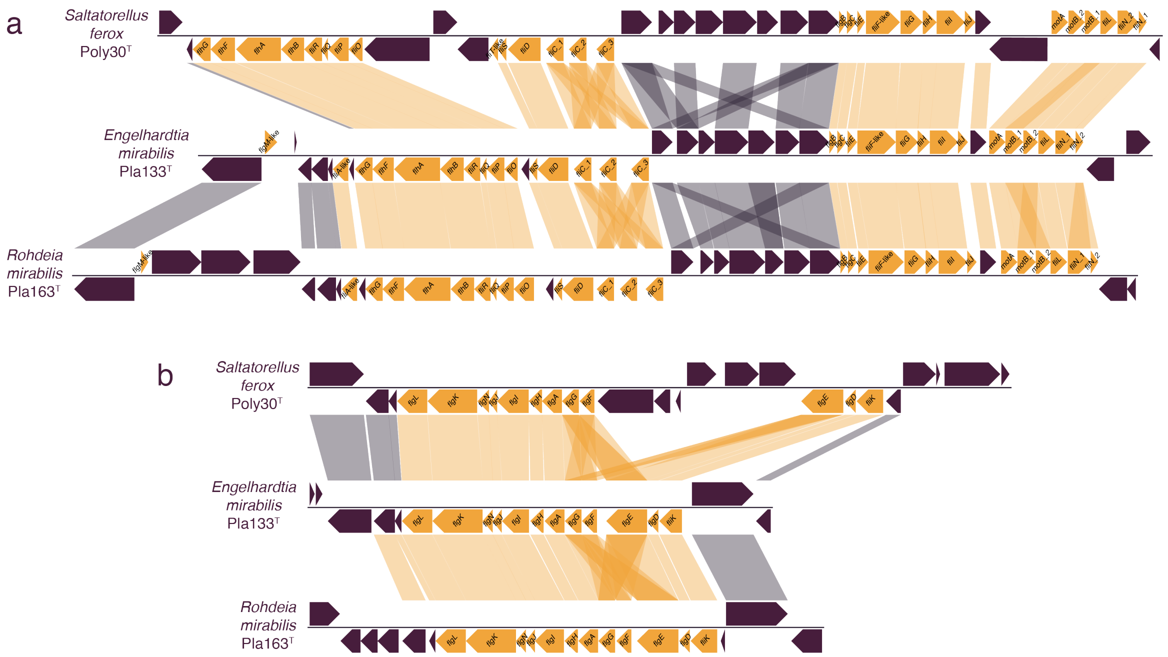

**Fig. S4. Genomic analysis of *Saltatorellota* locomotion: flagella.** Both panels (a and b) show two distinct locations on the chromosomes of all three *Saltatorellota* strains.

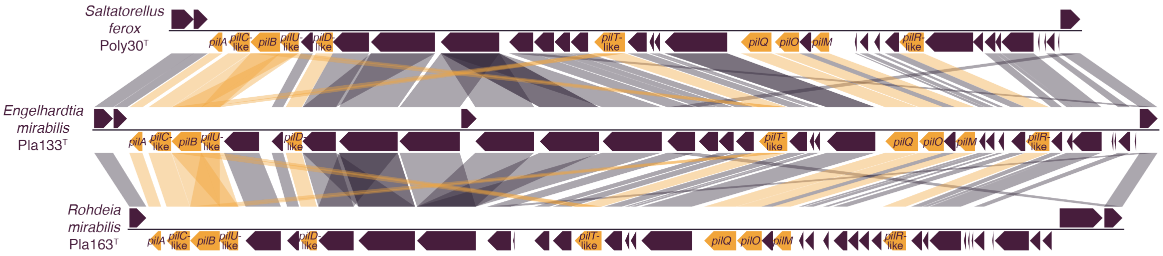

**Fig. S5. Genomic analysis of *Saltatorellota* locomotion: twitching motility related genes**. A gene locus in all three *Saltatorellota* genomes encodes most of the genes required for type 4 pili formation with potential usage in twitching motility.

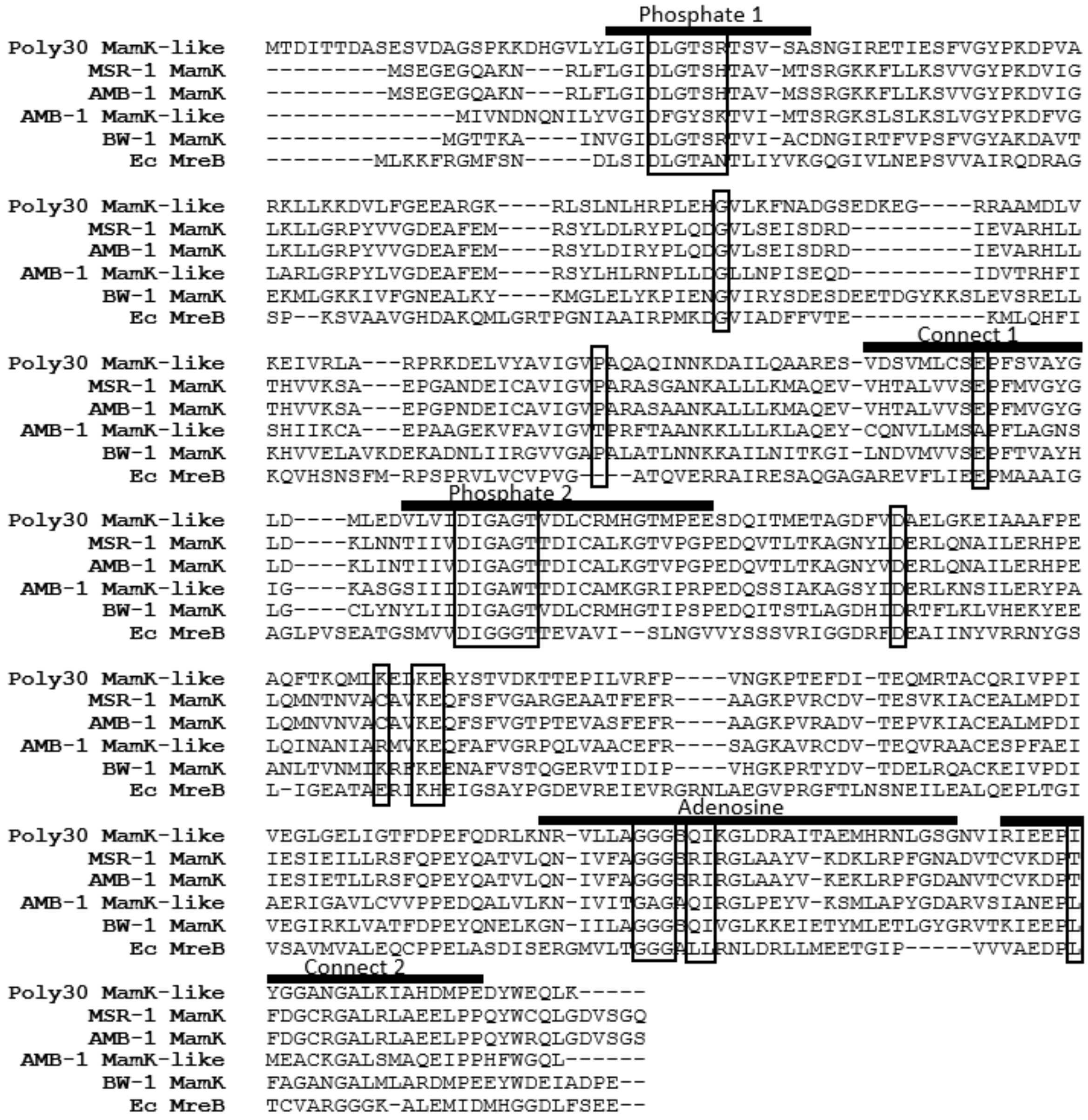

**Fig. S6. Alignment of MreB, MamK and MamK-like protein from *Saltatorellus ferox* Poly30$^T$.** MamK protein sequences from selected magnetotactic bacteria (MSR-1, *Magnetospirillum gryphiswaldense*; AMB-1, *Magnetospirillum magneticum*; BW-1, *Desulfamplus magnetovallimortis*), and MreB from *Escherichia coli* (Ec) were aligned with amino acid sequence of the MamK-like protein from *S. ferox* Poly30$^T$ applying MUSCLE alignment software packed in the Geneious Pro 8.1.3 software suite (https://www.geneious.com). Conserved residues necessary for the ATPase activity of MamK are boxed.

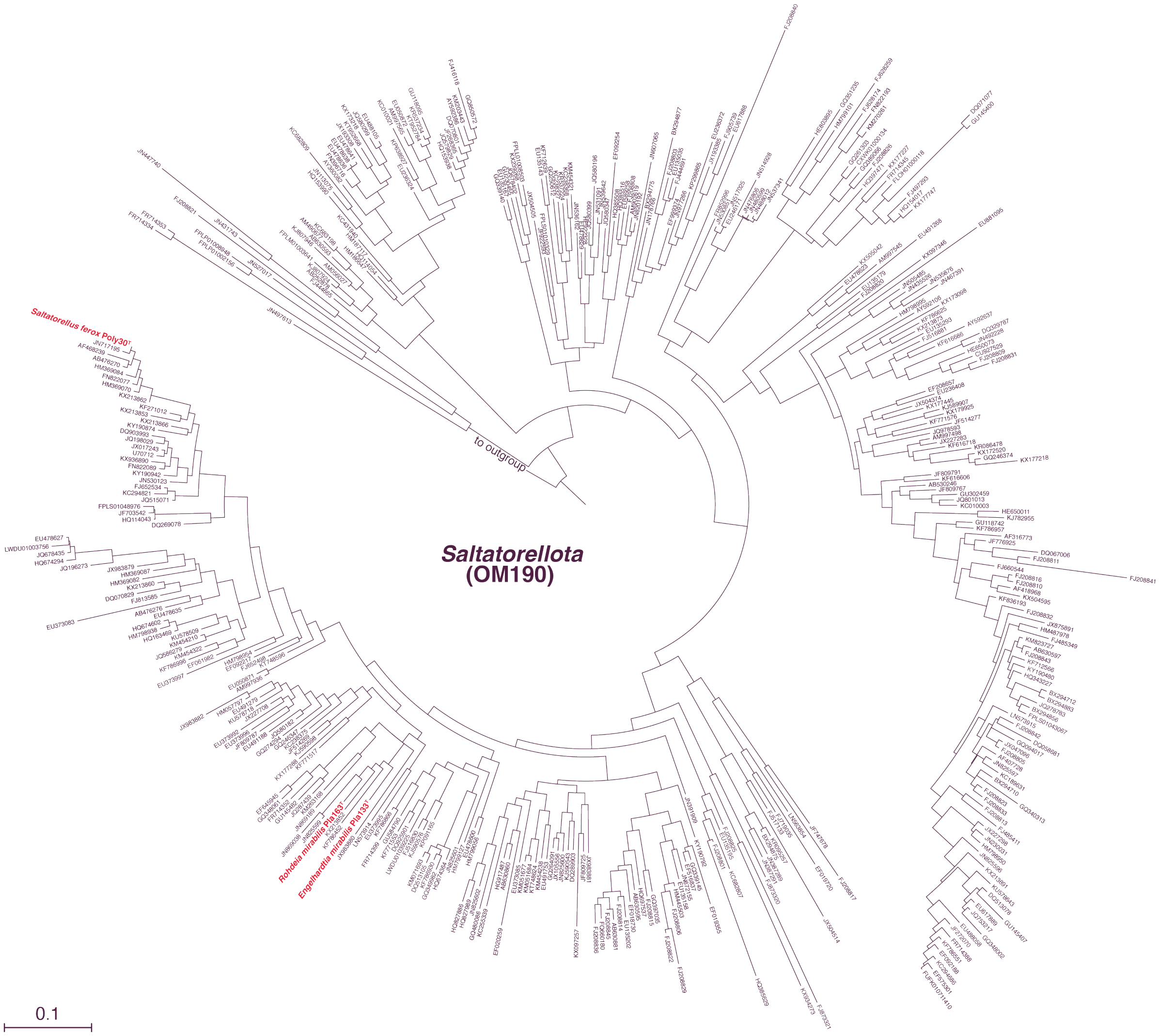

**Fig. S7. 16S rRNA sequence identity of the novel isolates.** Maximum likelihood phylogeny of 413 members of the SILVA taxonomy group OM190 and *Engelhardtia mirabilis* Pla133$^{T}$, *Rohdeia mirabilis* Pla163$^{T}$ and *Saltatorellus ferox* Poly30$^{T}$.

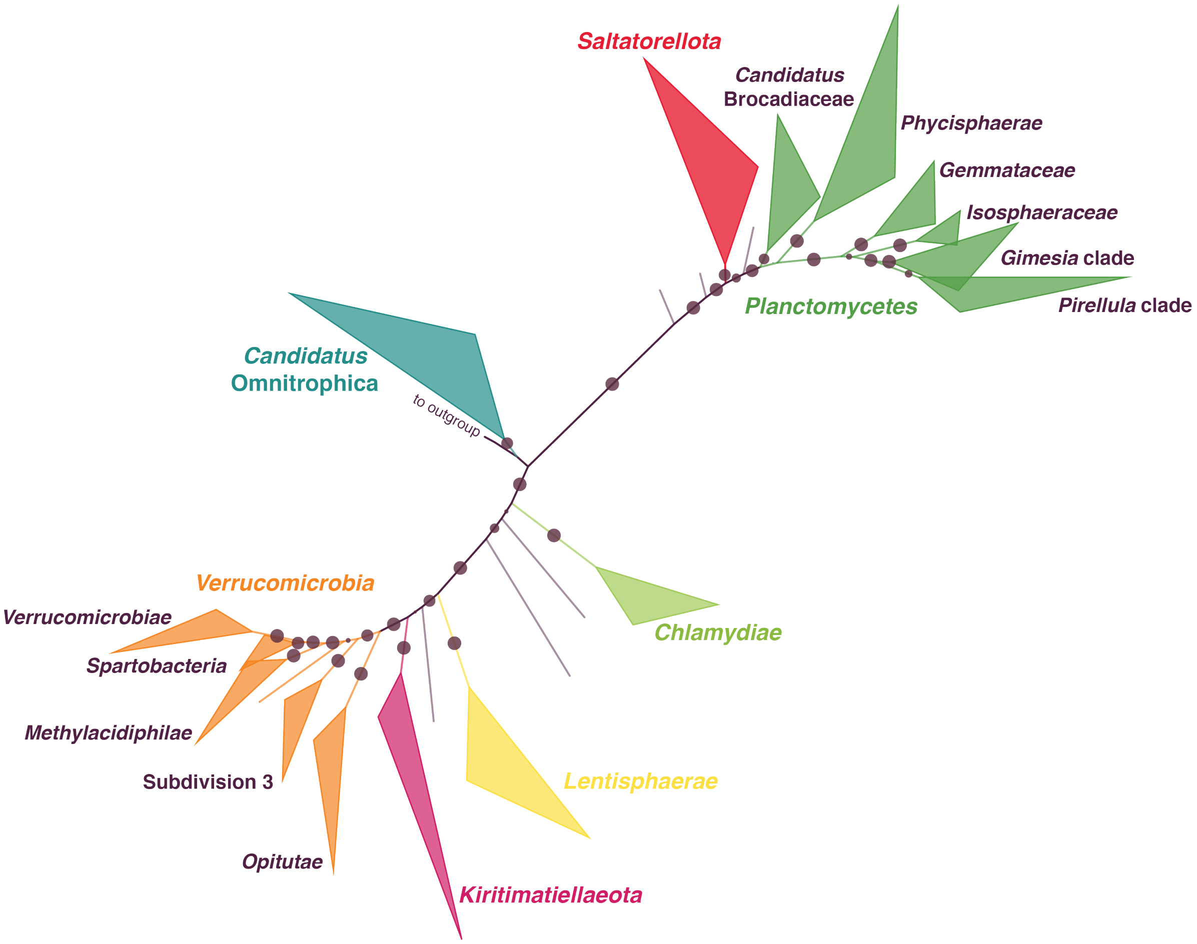

**Fig. S8. RpoB sequence-based phylogeny of the PVC superphylum.** Maximum likelihood phylogeny of 1631 RpoB protein sequences. The different colors indicate the different known phyla of the PVC superphylum. Subclades of the phyla *Verrucomicrobia* and *Planctomycetes* are indicated by equal coloring. The circles indicate reliability estimators (0.002 - 1) based on Shimodaira-Hasegawa testing (see Material and Methods).

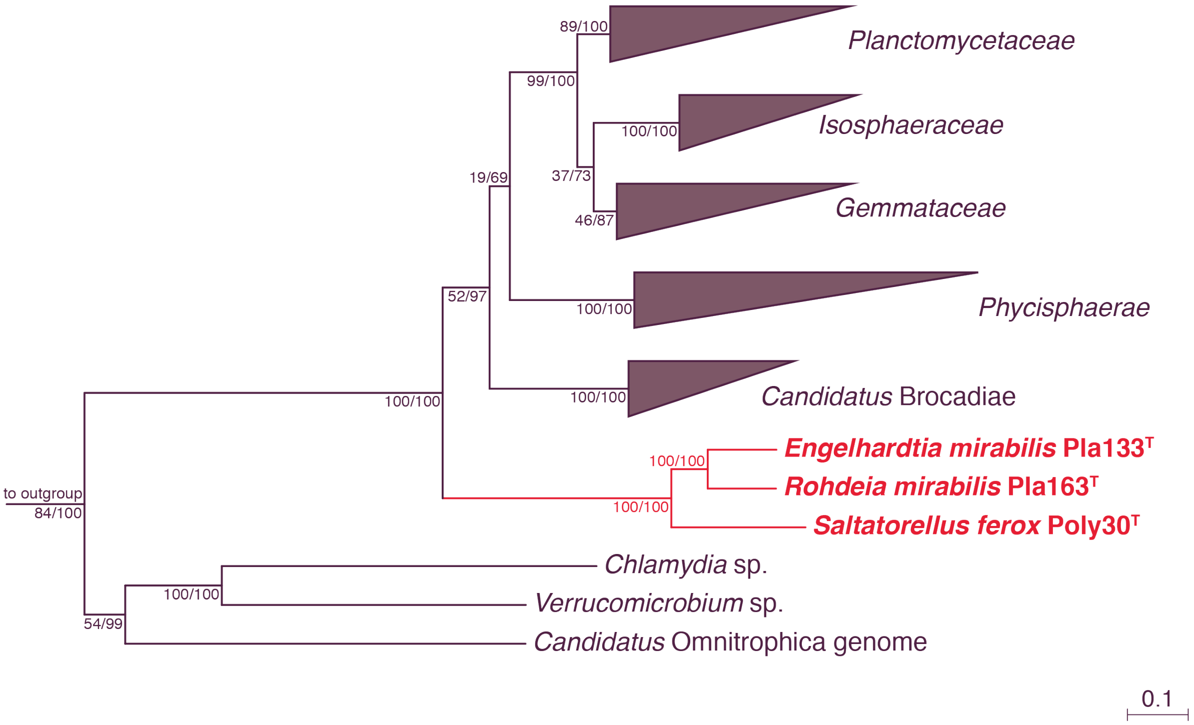

**Fig. S9. RpoB sequence-based phylogeny of the novel strains.** Maximum likelihood calculation for all described and proposed [1] planctomycetal genera and the three novel isolates Poly30$^{T}$, Pla133$^{T}$ and Pla163$^{T}$. The node values give a) the bootstrap values after 100 re-samplings and b) the posterior probability values after 300,000 generations (see Methods). The tree was rooted against three sequences from the phyla *Chlamydiae*, *Verrucomicrobia* and *Candidatus* Omnitrophica.

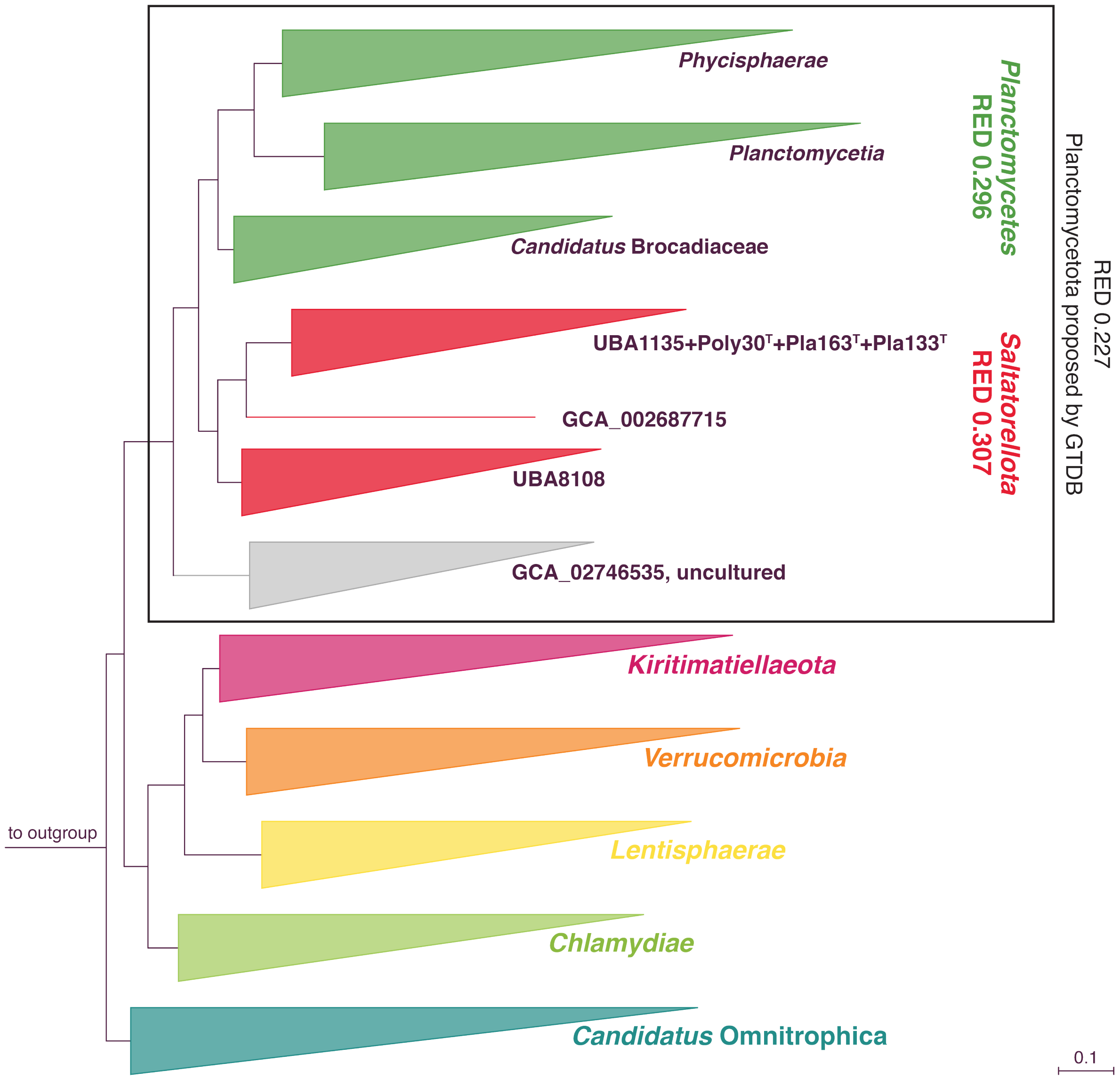

**Fig. S10. Phylogeny of the PVC superphylum.** Maximum likelihood phylogeny of a genome-based alignment from 120 concatenated protein marker genes [2] of 1675 genomes. The different colors indicate the different phyla of the PVC superphylum. RED values and GTDB taxonomy information are added. This tree is another depiction of the phylogeny shown in Fig. 7. For a more detailed phylogeny of the phylum *Saltatorellota* please refer to Fig. S11.

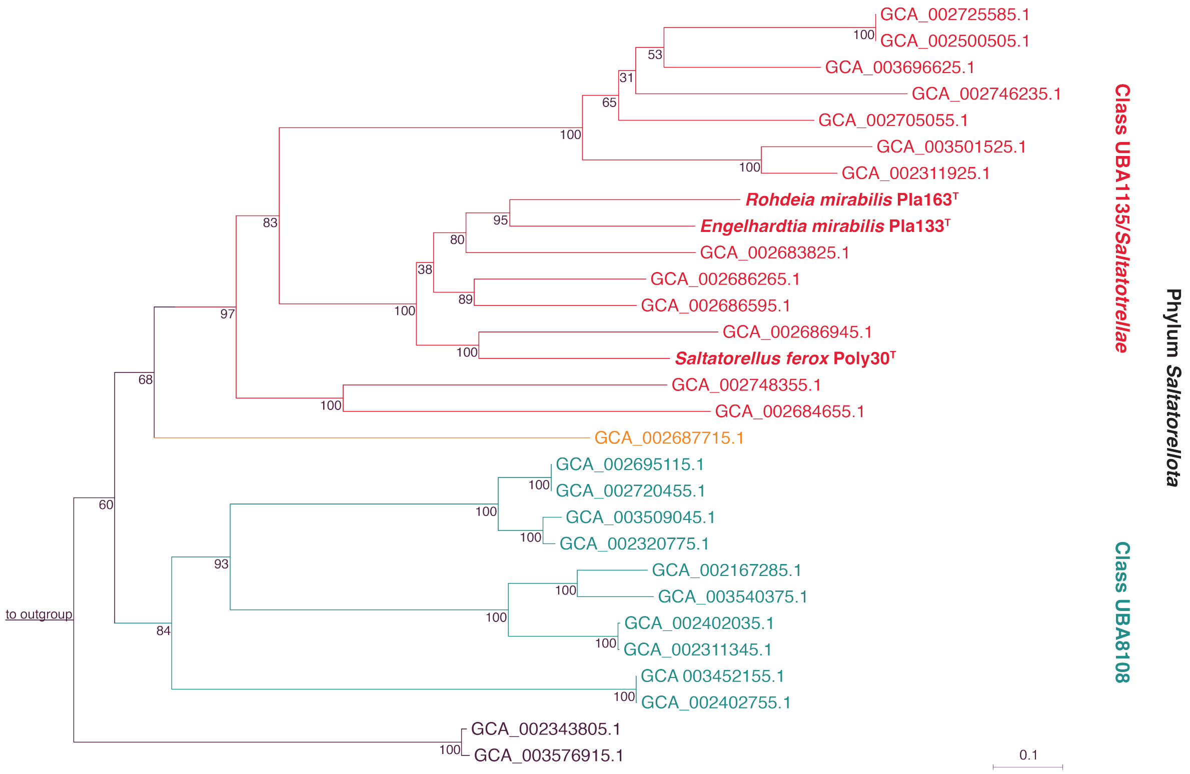

**Fig. S11. Multilocus sequence analysis of the phylum *Saltatorellota*.** Maximum likelihood phylogeny of well conserved parts of those seven protein sequences (1392 aa in total) present in all included genomes (novel isolates and MAGs). The node marks give bootstrap values after 100 re-samplings. The different colors point out the three GTDB taxa (class level) [2] included in the novel phylum *Saltatorellota*.

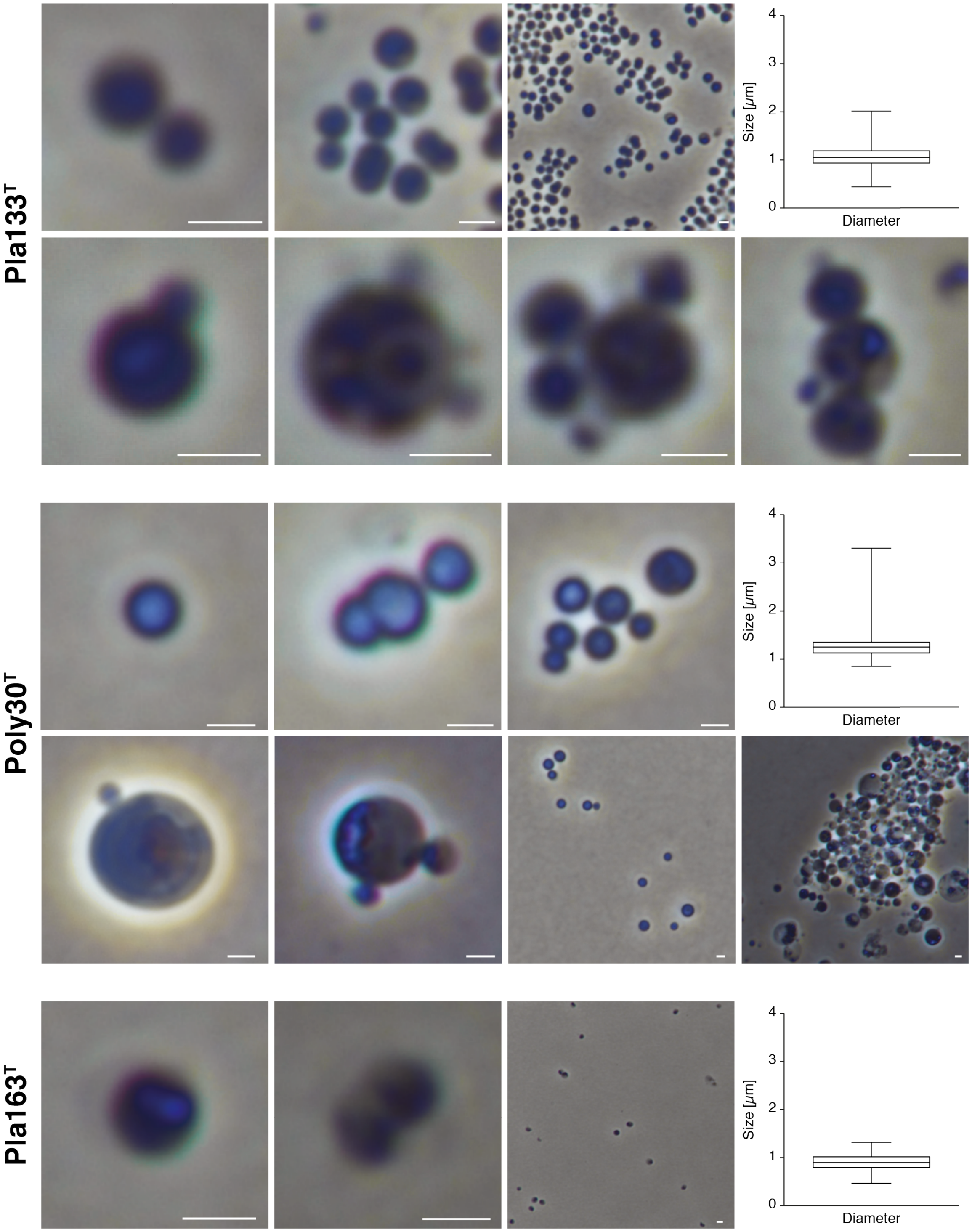

**Fig. S12. Cell size of *Saltatorellota* strains**. Phase contrast wide field light microscopic images of representative micrographs from *Engelhardtia mirabilis* Pla133$^T$, *Saltatorellus ferox* Poly30$^T$ and *Rohdeia mirabilis* Pla163$^T$. Average cell size is blotted. Scale bar 1 µm.

### Supplementary Tables

**Table S1. Detailed information on location, chemical parameters and abundance of *Saltatorellus ferox*, *Engelhardtia mirabilis* and *Rohdeia mirabilis* for each sampling point during the Baltic Sea cruise.** The abundance was calculated as relative abundance compared to all bacteria, based on 97% 16S rRNA gene identity. The attached-living fraction was obtained by filtration through a 3 µm pore size filter. The flow-through was filtered by a 0.2 µm pore size filter to obtain the free-living fraction. (see xlsx file)

**Table S2. Analysis of motility-associated genes.** For all identified candidate genes a Pfam and TIGRFAM analysis was carried Out as described in the Methods. For all positive hits, the locus tag of the hit is given. (see xlsx file)

**Table S3. Specifications for Fig. S4 and S5.** The annotations and gene names for all identified depicted genes are given. The identification was either done by Pfam/TIGRFAM hit or by InterProScan analysis when found in putative cluster. (see xlsx file)

**Table S4. Protein similarities.** Percentages of similarity between the MamK-like protein of the *Saltatorellus ferox* Poly30$^T$ compared to the selected magnetotactic bacteria *Magnetospirillum gryphiswaldense* MSR-1, *Magnetospirillum magneticum* AMB-1, *Magnetovibrio* MV-1 and strain BW-1.

| | **MamK$_{MSR-1}$** | **MamK$_{AMB-1}$** | **MamK-like$_{AMB-1}$** | **MamK$_{BW-1}$** |
|---|---|---|---|---|
| **MamK-like$_{Poly30}$** | 70.94% | 70.94% | 65.71% | 78.25% |

**Table S5. Minimal 16S rRNA gene sequence identity values within the given clades.** Values <75% indicate the inclusion of more than one phylum.

| **Possible taxa** | **Minimal 16S rRNA gene identity** |
|---|---|
| Planctomycetes + Poly30$^T$, Pla133$^T$, Pla163$^T$ | 71.8% |
| Poly30$^T$, Pla133$^T$, Pla163$^T$ | 89.0% |
| Planctomycetes | 71.8% |
| Poly30$^T$, Pla133$^T$, Pla163$^T$ | 89.0% |
| Candidatus Brocadiaceae | 87.0% |
| Phycisphaerae | 77.8% |
| Planctomycetia | 76.4% |

**Table S6. 16S rRNA sequence identity matrix of SILVA taxon OM190.** All identity values are given in percent. The grey-shaded strains are *Planctomycetes* for comparison. *Saltatorellota* strains Poly30$^T$, Pla133$^T$ and Pla163$^T$ are marked by red font. (see xlsx file)

**Table S7. Relative evolutionary divergence (RED) values within the given clades.** Values between 0.226 and 0.426 indicate described or potential phyla with two or more subordinate taxa. Values above this threshold indicate taxa with only one class. The green shaded taxon Planctomycetota as proposed by GTDB comprises 10 different taxonomic classes [2] including UBA1135, UBA8108, GCA-002687715. The yellow shaded phyla are as proposed by this study and the blue shade indicates taxonomic ranks within the phylum *Planctomycetes* that could be classified as separate phyla.

| **Possible phyla** | **RED values** |
|---|---|
| Planctomycetota (as proposed by GTDB) | 0.227 |
| *Saltatorellota* (Poly30$^T$, Pla133$^T$, Pla163$^T$, UBA1135, UBA8108, GCA-002687715) | 0.307 |
| *Planctomycetes* | 0.296 |
| *Candidatus* Brocadiaceae | 0.33 |
| Phycisphaerae | 0.393 |
| Planctomycetia | 0.454 |

### Supplementary Movies

**Movie S1.** Budding and binary fission in *Engelhardtia mirabilis* Pla133$^T$.

**Movie S2.** Cell fusion in *Engelhardtia mirabilis* Pla133$^T$.

**Movie S3.** Amoeba-like locomotion and shapeshifting in *Saltatorellus ferox* Poly30$^T$.

**Movie S4.** Detailed locomotion and shapeshifting in *Saltatorellus ferox* Poly30$^T$.

**Movie S5.** Locomotion in daughter cells of *Saltatorellus ferox* Poly30$^T$.

**Movie S6.** Shapeshifting in *Saltatorellus ferox* Poly30$^T$.

**Movie S7.** Trunk formation in *Saltatorellus ferox* Poly30$^T$.

**Movie S8.** Growing aggregate of *Saltatorellus ferox* Poly30$^T$.

**Movie S9.** Growing aggregate of *Saltatorellus ferox* Poly30$^T$ with single cells.

**Movie S10.** Trunk-forming aggregate of *Saltatorellus ferox* Poly30$^T$.
